# Mitigating Cellular Dysfunction by Addressing Contaminants in Synthetic circRNA

**DOI:** 10.1101/2024.09.17.613157

**Authors:** Ziwei Zhang, Weiyu Li, Dengwang Luo, Xiushuang Yuan, Li Yu, Daming Wang, Yuhong Cao

## Abstract

Synthetic circular RNA (circRNA) shows great potential in biomedical research and therapies, but impurities introduced during its synthesis can undermine circRNA efficacy. This study investigated the immunogenic potential of byproducts generated during its production. We found that trace amounts of double-stranded RNA in high molecular weight impurities, 5’ triphosphates of uncircularized RNA or small intron fragments, and hydrolyzed RNA fragments significantly impact the functionality of circRNA by activating innate immune responses through the sensory molecules involved in RNA sensing. To address this, we developed a novel multi-step purification process that employs enzymatic treatments and cellulose-based filtration to selectively remove these detrimental contaminants. This tailored approach minimizes cellular immune reactions and substantially improves circRNA yields with up to more than 10 times increase. Our findings underscore the critical impact of precise contaminant management in enhancing the expression and potential therapeutic utility of circRNA. This suggests a new direction for optimizing their production for both research and clinical applications.

## Introduction

Synthetic circular RNA (circRNA) is gaining prominence in biomedical research due to their unique covalently closed structure, which lacks 5’ or 3’ ends, protecting it from exonuclease degradation (1, 2). This structural stability enables sustained protein expression, giving circRNA significant advantages over traditional linear mRNA in therapeutic applications (1, 3–7). However, the potential immunogenicity of engineered circRNA remains a subject of debate, stemming from conflicting findings across multiple studies. Factors influencing these results include circRNA synthesis methods, presence of RNA modifications, purification techniques, and detection assays used to measure immune responses (8–12). Further research is needed to reconcile these differences and fully understand the immunogenic potential of engineered circRNA.

There are a few signaling pathways that can be activated by the presence of different forms of RNA (13). Toll-like receptors (TLRs) (14, 15) and cytoplasmic sensors like retinoic acid-inducible gene I (RIG-I) (16, 17) and melanoma differentiation-associated gene 5 (MDA5) (17, 18) play a critical role in responding to pathogen-associated molecular patterns in RNA, leading to immune activations that compromise the integrity and expression of RNA. Additionally, 2′-5′-oligoadenylate synthase (OAS) and RNA-dependent protein kinase R (PKR) are crucial in detecting RNA, with OAS activated by double-stranded RNA (dsRNA) leading to RNase L activation and PKR responding to a variety of RNA structures (19–21). These molecular sensors collectively initiate immune responses against RNA-based pathogens by detecting distinct RNA patterns and activating downstream signaling pathways (13, 22).

The intracellular expression efficiency and biosafety of circRNA is often compromised by RNA byproducts produced during synthesis. The immunogenic byproducts of circRNA arise during both the *in vitro* transcription (IVT) and RNA circulation processes (8). RNase R (RR) digestion is widely applied for the removal of linear byproducts in circRNA. However, highly structured RNAs, particularly those with double-stranded regions and stable secondary structures, can be resistant to degradation (22). Other commonly used methods, such as size-exclusion liquid chromatography (SEC), are inadequate for removing immunogenic contaminants from circRNA preparations, which can trigger innate immune responses in transfected cells, reducing protein expression, raising safety concerns and often resulting in low yields [8]. These limitations increase production costs and time, limiting the potential for large-scale manufacturing and widespread use of circRNA. The purification challenges underscore the need for an optimized purification strategy capable of effectively eliminating contaminants that activate immune responses.

In this study, we performed a comprehensive analysis of the immunogenicity of RNA byproducts during circRNA synthesis and identified four key contaminants: RNA precursors, hydrolyzed circRNA (nicked RNA), dsRNA, and RNA 5’ triphosphate. We demonstrated that even trace amounts of dsRNA can significantly stimulate cells and affect circRNA expression. In contrast, the immunostimulatory effects of small quantities of linear RNA and introns can be mitigated by phosphatase treatment, which removes the 5’ triphosphate groups recognized by innate immune sensors. This highlights the importance of thorough purification strategies to eliminate contaminating RNA species, as even minute amounts of impurities can confound the interpretation of circRNA immunogenicity and expression studies. To eliminate the immunogenicity in synthetic cricRNA, we developed a stepwise purification strategy that combines cellulose filtration with enzymatic digestion. This new purification approach not only mitigates immune responses but also enhances circRNA expression, achieving a significant increase in production efficiency compared to traditional methods. Furthermore, it significantly improves yields relative to conventional size-based purification techniques.

## Results

### Byproducts in synthetic circRNA significantly trigger innate immune responses

In the synthesis of circRNA, transcription from a DNA template is followed by the circularization of linear precursors through either direct ligation with T4 DNA/RNA ligase or self-splicing via group I/II autocatalytic introns (1, 26). The immunogenicity of circRNA can stem from the formation of dsRNA byproducts, uncircularized RNA (precursor), or nicked RNA molecules, all of which are potent triggers of cellular immune mechanisms.

To identify the immunogenic RNA byproducts in synthesis circRNA, we employed a circRNA encoding enhanced green fluorescent protein (circ-eGFP) and using coxsackievirus b3 as internal ribozyme entry site (IRES). Circ-eGFP was synthesized via group I intron-based method known for its high circulation efficiency (Fig. 1a). Capillary electrophoresis analysis of the unpurified circRNA preparation revealed four distinct peaks: the low molecular weight peaks for excised introns of the circularization process, a main circRNA peak that contained the desired circular RNA, a precursor peak, and a high molecular weight (HMW) peak that consisted of HMW byproducts such as dsRNA and other RNA aggregates (fig. S1a). To explore the immunogenicity of different byproducts, we employed SEC to fractionate the unpurified circ-eGFP into distinct components, including HMW, main, and intron fractions (Fig. 1b). Capillary electrophoresis confirmed that each fraction exhibited different profiles of impurities (Fig. 1c). The intron fraction contained introns and a small proportion of the main peak, whereas the main fraction contained circRNA, nicked RNA, and linear precursors. The HMW fraction primarily consisted of HMW byproducts such as dsRNA and other RNA aggregates.

**Fig. 1.**
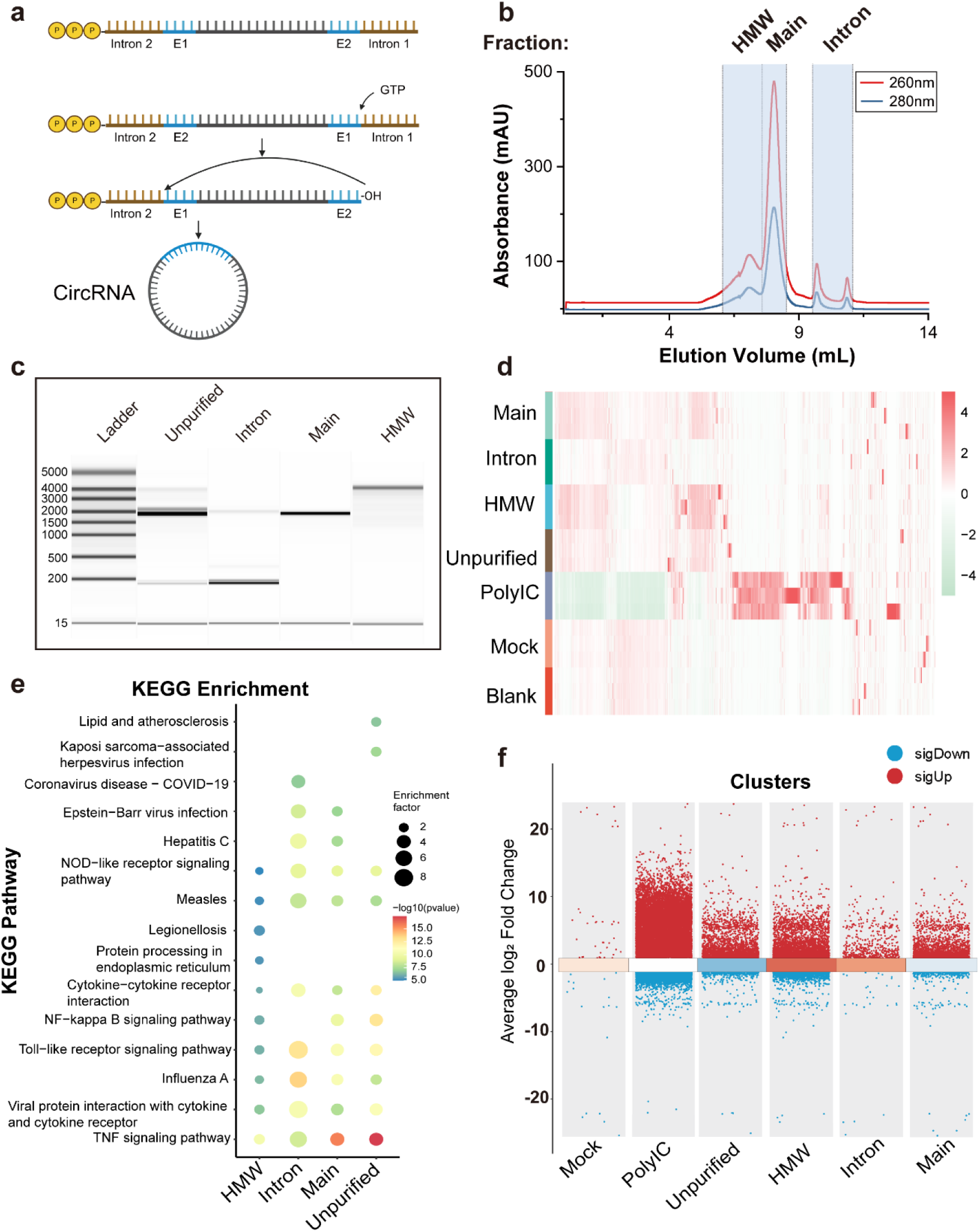
Innate immune activation by byproducts of IVT-synthesized circular RNA. **a,** Schematic representation of RNA circularization with group I autocatalytic introns (Created with BioRender.com). **b,** Size exclusion chromatography separation of unpurified circRNA. **c,** Capillary electrophoresis of different fractions, as described in **b**. **d,** Gene expression profiles of PMA-differentiated THP-1 cells transfected with different fractions, each group presented with three biological replicates. **e,** Multiple KEGG enrichment analysis of PMA differentiated THP-1 transfected with different fractions, with untreated cells serving as the control group. **f,** Multiple volcano plots illustrating transcriptomic differences between THP-1 cells transfected with distinct fractions. Untreated cells served as the control group. Each cluster represents the comparison of a specific fraction-transfected condition against the control.

To investigate the mechanisms underlying the immunogenicity of circRNA and its byproducts, we transfected unpurified circRNA, main fraction, HMW or intron fraction, into PMA-differentiated THP-1 cells to exam their immunogenicity in cells (Methods). Transcriptome analysis revealed that the intron-treated group showed relatively moderate differences, whereas the unpurified circRNA, main fraction and HMW fraction-treated cells exhibited marked changes in gene expression patterns (Fig. 1d-f).

Kyoto Encyclopedia of Genes and Genomes (KEGG) pathway enrichment analysis highlighted significant differences between the blank and different fractions, particularly in immune response and viral infection pathways. Key enriched pathways included TNF signaling, TLR signaling, NF-κB signaling, and cytokine-cytokine receptor interaction, which are essential for immune activation and inflammation. The unpurified cricRNA group exhibited the highest enrichment factors and most significant p-values. This suggested a strong association with these pathways in the main fraction under unpurified conditions. A similar profile can be observed in the main fraction group, likely due to the presence of linear precursors and nicked RNA. Intron and HMW fractions were transfected even in smaller amounts (Methods) also displayed remarkable enrichment in immune response and viral infection pathways. Notably, the intron fraction group did not involve the NF-κB signaling pathway, indicating that the introns lack typical functional motifs or structures required to engage immune receptors that trigger this pathway.

Exogenous RNAs, including single-stranded RNA (ssRNA) and dsRNA, are recognized by endosomal TLR and cytosolic RNA sensors (MDA5/RIG-I), triggering downstream signaling cascades that lead to the expression of inflammatory cytokines and interferons, as well as the activation of RNA degradation mechanisms mediated by PKR and OAS molecules (Fig. 2a) (13, 27). We identified potential candidate genes upregulated by unpurified circRNA in the context of cellular mechanisms for recognizing foreign RNA and the downstream innate immune response (Fig. 2b). The results suggested that unpurified cricRNA, main and HMW fractions contained byproducts had a significant upregulation of the RNA sensors, including TLR3, TLR7, TLR8, PKR, OSA, MDA5 and RIG-I, leading to the activation of downstream signaling pathways and triggering an inflammatory response by inducing cytokines, chemokines, and interferons. In contrast, the intron fraction displayed much lower activation of immune innate response compared to main and HMW fractions.

**Fig. 2.**
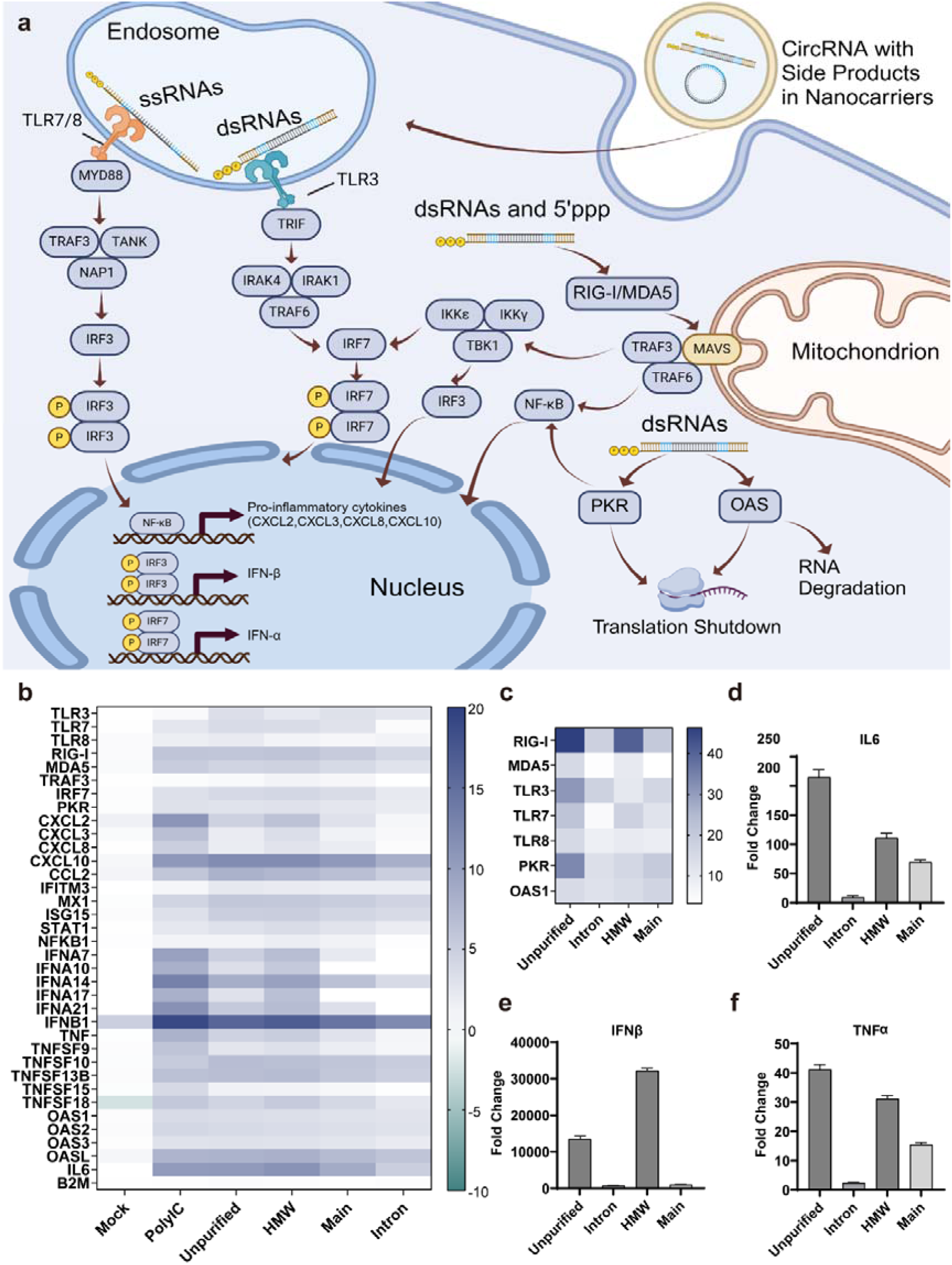
RNA sequencing revealed immunogenic potential of IVT-synthesized circRNA and byproducts. **a**, Cellular mechanisms for sensing foreign RNAs (Created with BioRender.com). **b,** Genes associated with innate immune response were identified as differentially expressed candidate mRNAs upon unpurified, intron, early and HMW transfection relative to untransfected cells, data presented as means of three biological replicates. B2M mRNA was used as housekeeping controls. **c,** Induction of RIG-I, TLR3/7/8, MAD5, PKR and OSA1 transcripts 6 hours post-transfection of PMA-differentiated THP-1 with unpurified, intron, main and HMW fractions relative to untransfected cells, Data presented as means of three biological replicates. B2M mRNA was used as housekeeping controls. **d-f,** Induction of IFNβ, TNFα, and IL6 transcripts 6 hours post-transfection of PMA-differentiated THP-1 with unpurified, intron, main and HMW relative to untransfected cells, B2M mRNA was used as housekeeping controls. Data presented as means ± SDs of three biological replicates.

Furthermore, quantitative real time-PCR (RT-qPCR) was employed to validate the innate immune response induced by different fractions (Fig. 2c-f). The unpurified circ-eGFP exhibited the high induction of sensory molecules involved in RNA sensing, such as RIG-I, MDA5, TLR3, TLR7, OAS, and PKR, suggesting the presence of a complex mixture of immunogenic RNA species, including dsRNA, 5’ triphosphates introns and precursors, and nicked RNA, as illustrated in fig. S2. Similarly, the HMW fraction induced substantial upregulation of these sensors, particularly the dsRNA sensors RIG-I and MDA5. The dsRNA aggregates in the HMW fraction may contribute to this enhanced immunogenicity. The main fraction exhibited moderate induction levels, potentially containing a mixture of linear RNA species (precursors with 5’ triphosphates and nicked RNA). In contrast, the intron fraction induced a markedly lower immune response, likely due to the presence of 5’ triphosphates or other structural features in the intron RNA that can be recognized by sensors like RIG-I and TLR sensors.

Regarding cytokine induction, the unpurified circ-eGFP and the HMW fraction triggered the highest expression levels of IFNβ, TNFα, and IL6. The potent induction of IFNβ by the HMW fraction can likely be attributed to the high levels of dsRNA (28). Conversely, the intron fraction exhibited the lowest cytokine activation, while the main fractions induced moderate cytokine production.

These results revealed the heterogeneous nature of unpurified circRNA. The unpurified circ-eGFP and HMW fraction were found to be the most immunogenic, indicating the presence of diverse RNA species that can activate multiple RNA sensors. In contrast, the intron fraction showed relatively low immunogenicity. Thus, effective purification methods may be necessary to obtain circRNA suitable for therapeutic applications.

### dsRNA removal is a critical step to improve circRNA performance

Our findings indicate that trace HMW impurities detrimentally impact circRNA functionality. These impurities pose a significant obstacle to achieving high-purity circRNA. Previous research has proven that microcrystalline cellulose (MCC) exhibits selective binding capacities for dsRNA versus ssRNA within an ethanol-HEPES buffer system (29). We employed MCC chromatography to investigate its efficacy in eliminating HMW impurities that adversely affect circRNA functionality and protein expression.

We collected both the flow-through peak and subsequent elution peaks, as detailed in Fig. 3A. The composition of these fractions was analyzed via capillary electrophoresis, revealing distinct distributions of RNA species; introns were primarily found in the flow-through and in early elution peaks, while HMW species were mainly present in the later elution peaks (fig. S3a-c, table S1). A lateral flow strip assay (LFSA) confirmed the absence of dsRNA in the early elution peaks, indicating that dsRNA was removed from these fractions (Fig. 3B) (30). Further protein expression tests conducted in HeLa cells assessed the functional implications of these purifications. The initial halves of the elution peaks under both equilibration conditions significantly enhanced protein expression and transfection efficiency, while later fractions showed diminished or no improvement in these parameters (Fig. 3c-e). Interestingly, the flow-through fraction with the highest intron content also demonstrated a notable increase in protein expression, further underscoring the negative impact of HMW impurities, which were strongly inverse-correlated with protein expression outcomes (R² = 0.84, fig. S3D)

**Fig. 3.**
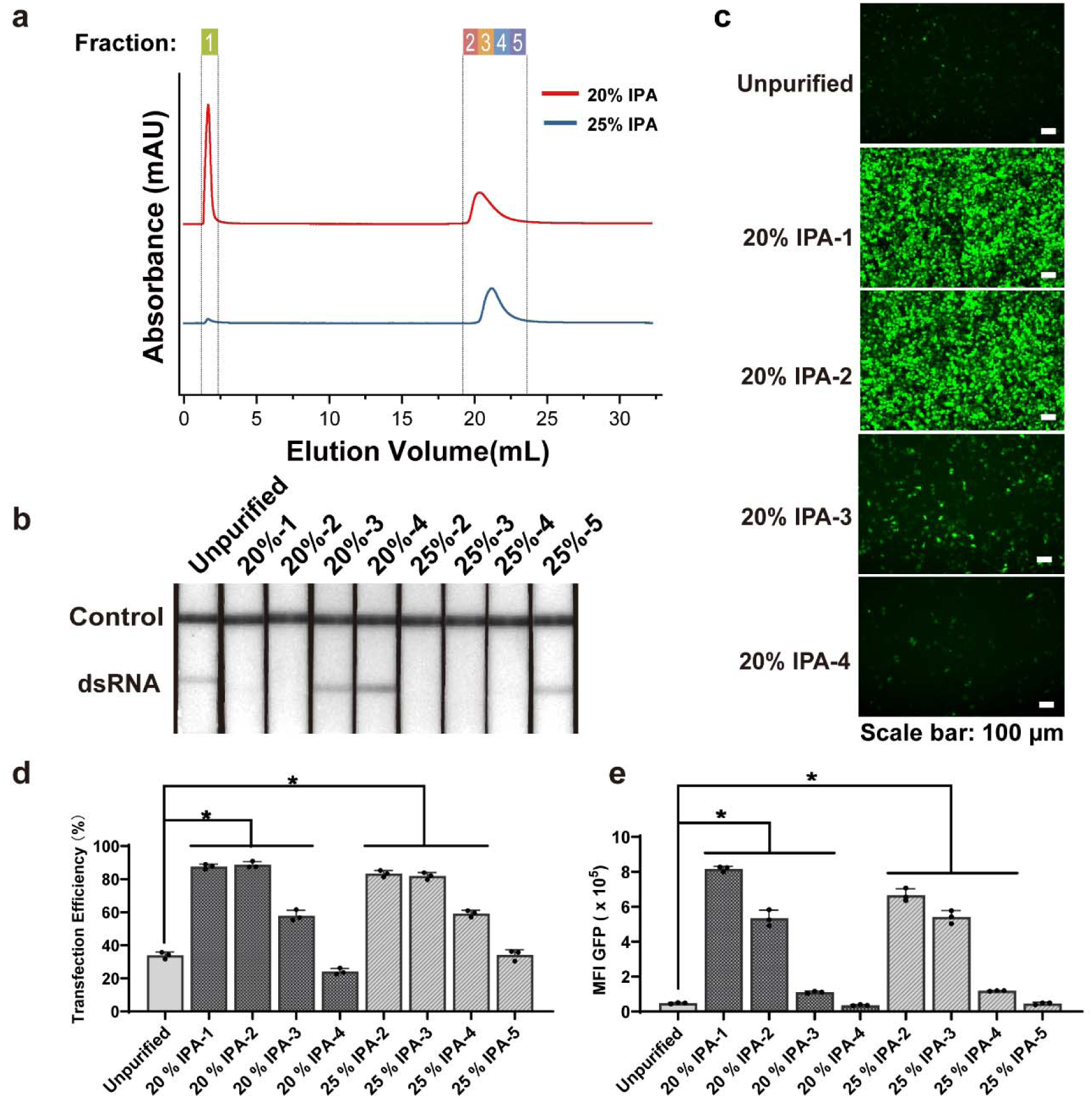
dsRNA removal significantly improves circRNA translatability. **a,** MCC chromatogram of unpurified circ-eGFP utilizing 20 % (red) and 25 % (blue) isopropanol (IPA)-HEPES equilibrium conditions. **b,** LFSA assessment of circRNA fractions as outlined in **a**. **c,** Representative cellular fluorescence image 24 hours post-transfection of HeLa cells with different fractions as outlined in **a** (scale bar: 100 μm). **d,** Translation efficiency and **e** mean fluorescence intensity (MFI) 24 hours post-transfection of HeLa cells with different fractions as outlined in **a**. Data presented as means ± SDs of three biological replicates, *p < 0.05, according to Student’s t-test.

### Enzymatic treatments are essential for improving circRNA purity and reducing immunogenicity

The removal of dsRNA can largely enhance the translatability of circ-eGFP. However, other contaminants such as linear species or introns can also stimulate immune responses and diminish circRNA expression. Therefore, we investigated enzymatic treatments to deal with immunogenic sources of circRNA.

We subjected the unpurified circ-eGFP to RR treatment, which selectively degrades linear RNA while leaving circRNA intact. As illustrated in fig. S1B, RR digestion effectively eliminated the majority of linear RNA, resulting in a circ-eGFP purity of approximately 89.8 %, with residual constituents including 6.2 % introns and 4.0 % HMW species. To evaluate the immunogenic potential of RR-treated circ-eGFP (RR+circ-eGFP), we isolated its HMW, main peak, and intron fractions using SEC for subsequent analysis (Fig. 4a). These fractions were tested in A549 cells to assess their innate immune responses. The results indicated that both RR+circ-eGFP and HMW fractions, which contained dsRNA, still triggered a potent innate immune response, albeit less so for the main fraction where linear contaminants were largely removed by RR (Fig. 4b & fig. S4a). A moderate immunogenicity can be observed regarding intron fraction.

**Fig. 4.**
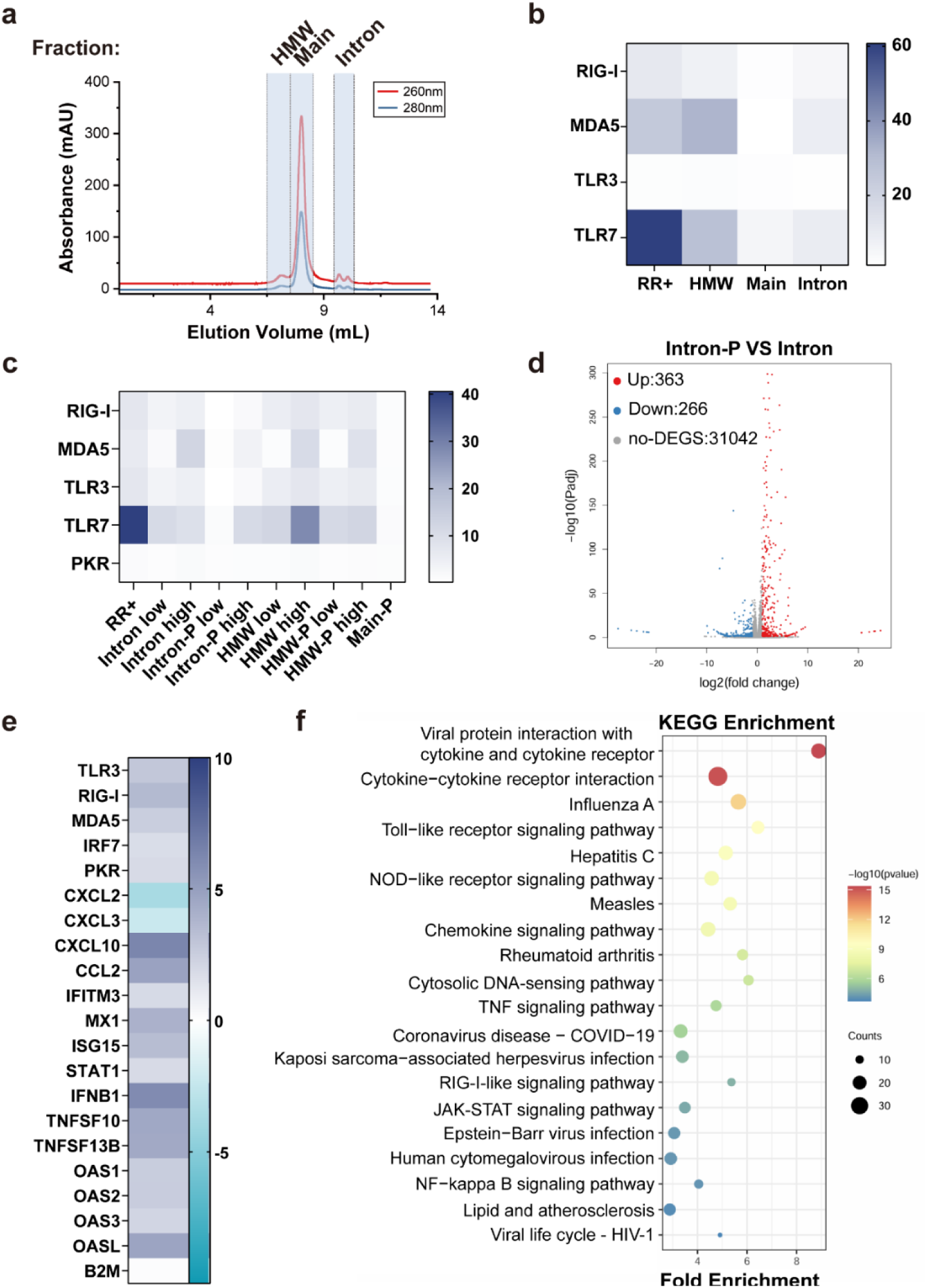
RNase R and Phosphatase to Purify IVT-produced circRNA. **a,** Size exclusion chromatography separation of RR+circ-eGFP. **b,** Induction of RIG-I, TLR3, TLR7, and MDA5 transcripts 6 hours after transfection of A549 cells with the 40 ng of intron, HMW or 200 ng of main fraction of SEC separated RR+circ-eGFP, data presented as means of three biological replicates. B2M mRNA was used as housekeeping controls. **c,** Induction of RIG-I, TLR3, TLR7, and MDA5 transcripts 6 hours after transfection of A549 cells with the 40 ng or 200 ng of intron and HMW, before and after phosphatase treatment (intron-P, HMW-P), or 200 ng of main fraction after phosphatase treatment (main-P), data presented as means ± SDs of three biological replicates. B2M mRNA was used as housekeeping controls. **d,** Volcano plots show the difference in transcriptome between THP-1 cells transfected with intron-P and intron. **e,** Genes associated with innate immune response were identified as differentially expressed candidate mRNA in intron-transfected cells relative to intron-P-treated cells, data presented as means of three biological replicates. B2M mRNA was used as housekeeping controls. **f,** KEGG enrichment analysis of PMA differentiated THP-1 transfected with intron relative to intron-P-treated cells.

To specifically address the immunogenicity associated with intron and HMW fractions, we treated intron, main and HMW fractions with phosphatase to remove 5’ triphosphate groups, known triggers for immune sensors such as RIG-I. Transfection into A549 cells with different dosage of HMW and intron showed that while phosphatase treatment reduced the immunogenicity of intron-derived impurities, it did little to mitigate the immune response elicited by HMW fractions, which primarily contain dsRNA (Fig. 4c & S4b). The main fraction, after phosphatase digestion, also displayed low immunogenicity. This can be attributed to the removal of the remaining 5’ triphosphates.

We used PMA-differentiated THP-1 cells to further explore the immunogenicity of the intron fraction after phosphatase treatment. Transcriptome analysis showed that 363 genes were upregulated, and 266 genes were downregulated in response to the intron fractions (Fig. 4d & e). KEGG and GO (Fig. 4f & S5a) analyses revealed that these differentially expressed genes were mainly involved in the innate immune response. To confirm these findings, we carried out RT-qPCR, which showed that the mRNA levels of RNA molecular sensors and cytokines decreased after phosphatase treatment (fig. S5b & c).

### Combined strategy removes immunogenic byproducts and yields high-purity circRNA

We have identified various immunogenic sources in IVT-synthesized circRNA, including dsRNA in HMW fractions, 5’ triphosphate introns, precursors, and nicked RNA. To obtain high-purity and low-immunogenicity circRNA, it is crucial to eliminate these impurities. Linear species can be removed through RR treatment, while the immunogenicity of 5’ triphosphates in introns and trace remaining precursors can be significantly reduced by phosphatase treatment. Additionally, dsRNA in HMW fractions can be effectively removed using cellulose chromatography.

To confirm that the contaminants are the major source of immunogenicity, we refined our purification protocol by integrating MCC chromatography with enzymatic treatments. This approach aimed to achieve even higher purity levels of circRNA while mitigating immunogenic responses. We purified RR+circ-eGFP using MCC (Fig. 5a), resulting in the isolation of an early-eluting peak, which was subsequently treated with phosphatase (RR+MCC-P). Capillary electrophoresis and LFSA results confirmed the effective removal of dsRNA from RR+MCC-P circ-eGFP, with only a minor fraction of introns (0.8 %) and HMW components (2.1 %) remaining (Fig. 5b & fig. S6a). An ultra-pure circ-eGFP was obtained by conventional purification method that involves SEC and enzymatic treatments was used as a positive control (fig. S6b, Methods, (8, 10)).

**Fig. 5.**
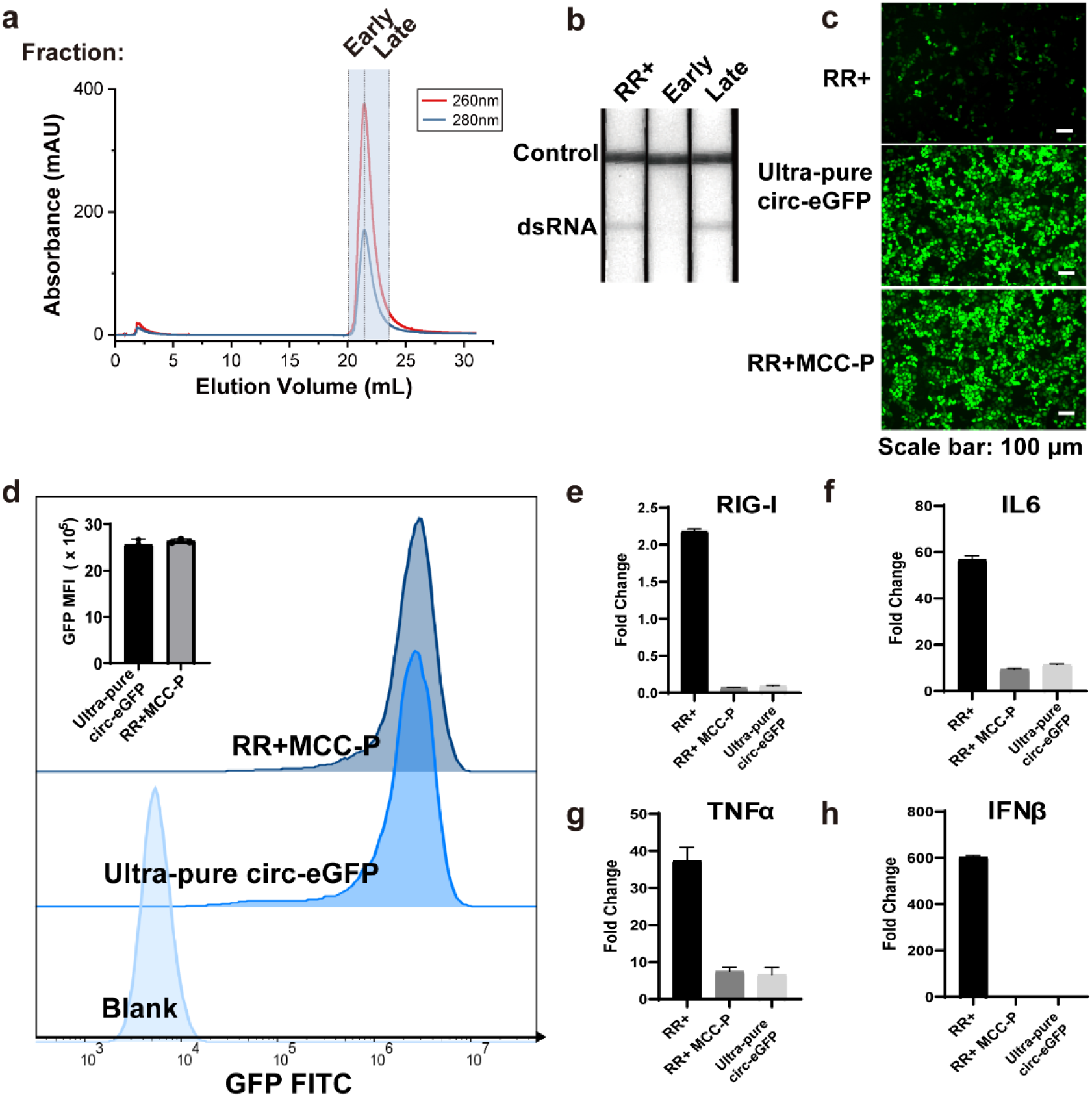
Enhancement of Protein Expression and Reduction of Immunogenicity via Combination of MCC Chromatography and Enzymatic treatment. **a,** MCC chromatogram of RR+ circ-eGFP under 25 % (blue) IPA HEPES equilibrium conditions. **b,** LSFA assessment of RR+, early and late peaks as detailed in **a**. **c,** Representative cellular fluorescence image (scale bar: 100 μm) captured 24 hours post-transfection of Hela cells with RR+, ultra-pure or RR+MCC-P circ-eGFP. **d,** Protein expression levels observed 24 hours post-transfection of HeLa cells with ultra-pure and RR+MCC-P circ-eGFP. Data presented as means ± SD of three biological replicates. **e-h,** Induction of RIG-I, IFNβ, TNFα, and IL6 transcripts at 6 hours post-transfection of 0.2 million A549 cells with 200 ng of RR+, RR+MCC-P, and ultra-pure circ-eGFP. Data presented as means ± SDs of three biological replicates.

RR+MCC-P circ-eGFP demonstrated protein expression levels that were comparable to those achieved by conventional treatments (Fig. 5c & d). To assess the immunogenic potential of these purified circRNA variants, we transfected A549 cells with both circ-eGFP under different purification conditions. RT-qPCR analysis revealed a significant reduction in the immunogenicity of circ-eGFP treated with RR+MCC-P, comparable to that observed with ultra-pure circ-eGFP (Fig. 5e-h). Notably, the mRNA expression levels of key innate immune sensors and cytokines, such as RIG-I, IFNβ, TNFα, and IL6, induced by RR+MCC-P were consistent with those induced by ultra-pure circRNA, confirming the effectiveness of our combined purification strategy in reducing immunogenic responses.

Despite the effective removal of dsRNA, a trace proportion of non-dsRNA HMW impurities remained in the purified circRNA. To further evaluate the influence of the remaining HMW, we conducted differential expression analysis using PMA-differentiated THP-1 cells. Compared to cells treated with HMW impurities after MCC treatment (HMW-MCC), 2762 genes were upregulated, and 1619 genes were downregulated in the group treated with HMW impurities before MCC treatment (Fig. 6a). Differential genes in the context of immune response were identified (Fig. 6b). KEGG enrichment and GO analysis (Fig. 6c & fig. S7) revealed that the major differences were related to immune response pathways, such as TNF signaling, cytokine-cytokine interaction, and TLR signaling. Thus, the removal of dsRNA can significantly alleviate immune responses, the remaining impurities, which is persistent to RR digestion, might be the dimmer form of circ-eGFP.

**Fig. 6.**
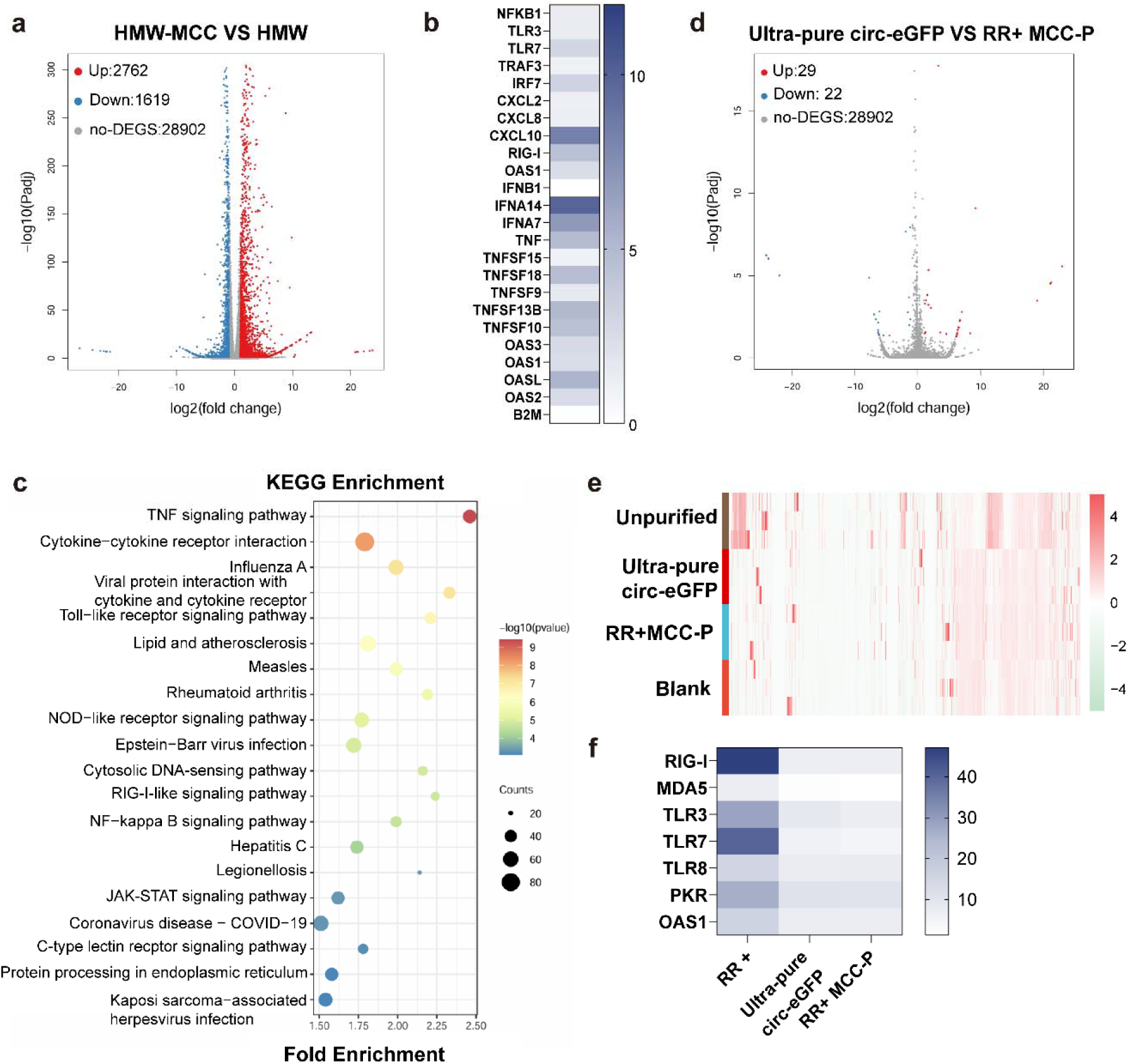
RNA sequencing revealed low immunogenicity of circRNA purified using enzymatic treatment and MCC chromatography. **a,** The volcano plot shows the difference in transcriptome between THP-1 cells transfected with HMW relative to the HMW-MCC treated cells. **b,** Genes associated with innate immune response were identified as differentially expressed candidate mRNAs upon HMW transfection relative to HMW MCC treated cells, data presented as means of three biological replicates. B2M mRNA were used as housekeeping controls. **c,** KEGG enrichment analysis of PMA differentiated THP-1 transfected with HMW, MCC, and HMW. **d,** The volcano plot shows the difference in transcriptome between THP-1 cells transfected with ultra-pure and RR+MCC-P circ-eGFP. **e,** Gene expression profiles of PMA-differentiated THP-1 cells transfected with ultra-pure, RR+MCC-P, and unpurified circ-eGFP, data presented as means of three biological replicates. **f,** Induction of RIG-I, TLR3, TLR7, PKR, OSA1 and MDA5 transcripts 6 hours after transfection of PMA-differentiated THP-1 cells transfected with ultra-pure, RR+MCC-P, and unpurified circ-eGFP, data presented as means of three biological replicates.

To comprehensively compare the immunogenicity of unpurified, ultra-pure, and RR+MCC-P circ-eGFP, we performed RNA sequencing, and the results are illustrated in Fig. 6d & e. Compared to unpurified circRNA, few differentially expressed genes were observed in the ultra-pure circRNA and RR+MCC-P groups relative to the untreated group. As expected, the ultra-pure circRNA and RR+MCC-P treated groups displayed similar transcription profiles. Further RT-qPCR results also confirmed the low immunogenicity of RR+MCC-Q (Fig. 6g& S8). These results validate the strategic integration of MCC chromatography and enzymatic treatments in our purification protocol, demonstrating that such a combined approach can efficiently yield circRNA preparations of high purity with significantly reduced immunogenicity.

### WMC chromatography efficiently purifies circRNA and minimizes innate immune response

Our findings have shown that IVT circRNA can be effectively purified by stepwise removal of immunogenic impurities, resulting in circRNA with high protein expression and low immunogenicity, comparable to the SEC method. To further improve the purification efficiency, we introduced wood-derived macroporous cellulose (WMC), a natural porous material, for the removal of dsRNA. We have previously demonstrated that the porous nature of WMC allows for efficient removal of dsRNA from circRNA of varying lengths while maintaining a high recovery rate of the target RNA (31). The optimized purification process consists of three key steps: (1) RR treatment to eliminate precursor RNA, (2) WMC chromatography for the removal of residual dsRNA, and (3) phosphatase treatment to eliminate potentially remaining phosphate groups (Fig. 7a).

**Fig. 7.**
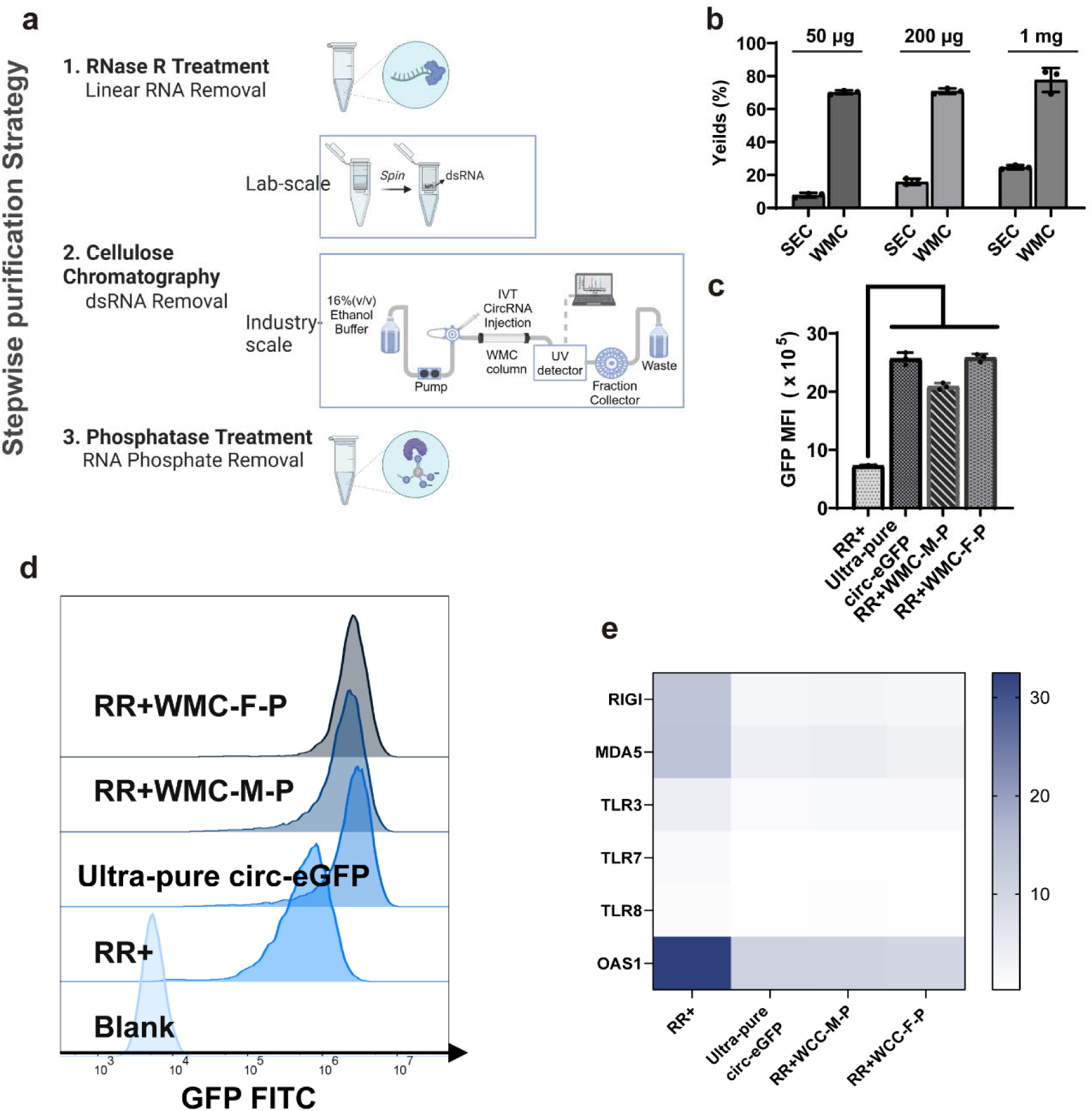
Optimized WMC chromatography to efficiently purify circRNA. **a,** Schematic representation of stepwise purification of circRNA (Created with BioRender.com). **b,** Recovery rate of RR+SEC, RR+WMC-M and RR+WMC-F chromatography Data presented as means ± SDs (n = 3). **c,** Protein expression level 24 hours post-transfection of HeLa cells with RR+, ultra-pure, RR+WMC-M-P and RR+WMC-F-P circ-eGFP. Data presented as means ± SDs of three biological replicates, *p < 0.05, according to Student’s t-test. **d,** MFI 24 hours post-transfection of HeLa cells with RR+, ultra-pure, RR+WMC-M-P and RR+WMC-F-P circ-eGFP. **e,** Induction of RIG-I, TLR3, TLR7, OSA1, PKR, IFN-β, TNF-α and IL-6 transcripts 6 hours after transfection of PMA-differentiated THP-1 cells with RR+, ultra-pure and RR+WMC-F-P circ-eGFP, data presented as means of three biological replicates.

We evaluated the recovery rate of WMC chromatography using different amounts of circ-eGFP (50 μg, 200 μg, and 1 mg). For 50 μg of circRNA, a micro-spin column with 5.3 mg of WMC was used (RR+WMC-M), while for 200 μg or 1 mg, a fast protein liquid chromatography (FPLC) system with 0.25 or 1.25 g of WMC was employed (RR+WMC-F), respectively (fig. S9a). The results demonstrated that WMC could completely remove dsRNA from circRNA across all tested RNA amounts (fig. S9b). Notably, WMC chromatography exhibited a remarkably higher recovery rate (> 70 %) compared to the conventional SEC method (RR+SEC), particularly for the 50 μg and 200 μg conditions (Fig. 7B).

The SEC, WMC-M, and WMC-F purified circ-eGFP samples were then subjected to phosphatase digestion to obtain the final products, designated as ultra-pure, RR+WMC-M-P, and RR+WMC-F-P, respectively. Capillary electrophoresis confirmed that both WMC-based methods achieved a purity of approximately 95 % (fig. S5c & d). Transfection circRNA into HeLa cells revealed that all WMC-based methods maintained high translation efficiency, identical to that achieved with ultra-pure circ-eGFP (Fig. 7c & d). Transcriptional analysis of innate immune receptors and cytokines indicated that WMC-purified circRNA elicited significantly reduced immunogenic responses, similar to those observed in ultra-pure samples (Fig. 7e & fig. S10). Overall, the stepwise byproduct removal strategy using WMC represents an efficient purification tool to produce high-yield, high-purity, and low-immunogenicity circRNA for therapeutic applications.

## Discussion

This study aimed to address the challenge of immunogenic impurities in synthetic circRNA, which can undermine their therapeutic potential. Comprehensive analysis revealed that contaminants such as dsRNA, RNA precursors, nicked RNA, and RNA phosphates significantly impact the immunogenicity and functionality of circRNA. These contaminants activate innate immune responses via RNA sensors such as PKR, OAS, TLR3, TLR7, TLR8, MDA5, and RIG-I, leading to reduced protein expression and potential cellular dysfunction.

The presence of dsRNA and other RNA contaminants in unpurified circRNA preparations was shown to significantly upregulate RNA sensors and inflammatory cytokines, particularly through pathways such as NF-κB, TNF, and TLR signaling pathways. This underscores the critical need for effective purification to ensure the biosafety and functionality of circRNA. Our work identified that even trace amounts of these immunogenic byproducts can activate immune responses and hinder circRNA expression.

To address these issues, the study developed a novel purification strategy involving a multi-step process combining enzymatic treatments and cellulose-based filtration. This method effectively removes the identified immunogenic impurities, thereby reducing the activation of immune responses and significantly improving the yield and purity of circRNA. The use of MCC or WMC chromatography was particularly effective in selectively binding dsRNA, further reducing immunogenicity. This novel purification approach resulted in highly pure circRNA with significantly reduced immunogenicity. The improved production efficiency achieved by this purification approach is approximately tenfold higher than traditional size-based methods, highlighting its potential for large-scale manufacturing.

The findings of this study have important implications for the production and application of circRNA in biomedical research and therapies. The ability to effectively purify circRNA and reduce immunogenic contaminants opens new possibilities for their use in preventing and treating diseases. Moreover, the integration of advanced purification techniques such as WMC chromatography with traditional methods could enhance the scalability and cost-effectiveness of circRNA production. Continued investigation into the sources and impacts of RNA impurities will be essential to refine purification processes and improve the overall quality and safety of synthetic circRNA.

Future efforts should prioritize refining and simplifying the multi-step purification process to facilitate broader industrial application. Developing a user-friendly, single-step purification method would be ideal for large-scale manufacturing, significantly enhancing its industrial applicability. To achieve this, minimizing undesired byproducts in crude products is essential. Optimizing T7 polymerase to reduce dsRNA byproducts at the transcription source can alleviate downstream purification burdens, effectively addressing primary immunogenicity issues and improving both safety and efficacy of the final product. Additionally, enhancing the efficiency of circRNA circularization will reduce linear RNA byproducts, further streamlining the purification process and improving overall yield and purity. These strategic refinements are crucial for the scalable, cost-effective production of high-quality circRNA, which is vital for their successful clinical and industrial applications.

In conclusion, this study provides a comprehensive approach to mitigating the immunogenicity of synthetic circRNA, paving the way for their safe and effective use in therapeutic applications. The novel purification strategy developed here represents a significant advancement in circRNA research, offering a robust solution to one of the key challenges in the field. The stepwise purification strategy developed in this study effectively addresses the challenge of immunogenic impurities, paving the way for safer and more effective circRNA-based therapeutics. By minimizing the activation of innate immune responses, this approach has the potential to enhance the clinical translation of circRNA technology.

## Materials and Methods

### Cell Lines

Human cervical carcinoma cell lines (HeLa), A549 and Human monocytic cell line THP-1 were obtained from the Cell Resource Center, Peking Union Medical College (which is the head quarter of National Science & Technology Infrastructure-National BioMedical Cell-Line Resource, NSTI-BMCR). HeLa and A549 were cultured at 37 °C and 5 % CO_2_ in Dulbecco’s Modified Eagle’s Medium (4500 mg/L glucose) supplemented with 10 % (v/v) heat-inactivated fetal bovine serum (hi-FCS, GIBCO) and 1 % (v/v) penicillin/streptomycin (GIBCO). THP-1 was maintained in RPMI 1640 GlutaMAX™ medium supplemented with 10 % (v/v) hi-FCS (FCS) and a 1 % (v/v) mixture of penicillin and streptomycin in a humidified incubator containing 5 % CO_2_ at 37 °C.

### circRNA in vitro synthesis

circRNA precursors were synthesized by T7 High Yield RNA Transcription Kit (Vazyme, China) according to manufacturer’s protocols. After transcription, reactions were treated with DNase I (Vazyme, China) for 15 min. After DNase treatment, circRNA was incubated from reactions with 8M LiCl. GTP was added to a final concentration of 2 mM along with a buffer including magnesium (50 mM Tris-HCl, 10 mM MgCl_2_, 1 mM DTT, pH = 7.5; New England Biolabs) into circRNA solution. Reactions was then heated to 55 for 8 min. After circularization, 20 U RNase R and 10× RNase R buffer (Epicentre) was added according to manufacturer’s protocols and was heated to 37 for 20 min. After RNase R digestion, circRNA was collected using LiCl precipitation. Then added 1 U/μg Quick CIP (New England Biolabs) and 10× rCurSmart into circRNA and heated 37 for 20 min.

### FPLC based purification of circRNA

450 mg of MCC was added into 1 mL column. The column was then connected to FPLC system and washed with chromatography buffer (10 mM HEPES,0.1 mM EDTA, 125 mM NACl and 25 % or 30 % (v/v) isopropanol) at a flow rate of 1 mL/min for 20 column volume. 100 μg of circRNA was loaded into the column. After purification, the isopropanol-containing eluent was precipitated with LiCl for recovery. After 250 mg of WMC was added into the 1 mL column, the WMC-FPLC based purification followed the same process with different chromatography buffer (10 mM HEPES, 0.1 mM EDTA, 125 mM NaCl, and 16 % (v/v) ethanol, pH = 7.2).

### Spin column based purification of circRNA

5.3 mg of WMC was filled into a spin column and washed using 500 μL elution buffer (10 mM HEPES, 0.1 mm EDTA, 125 mm NaCl, and 16 % (v/v) ethanol, pH = 7.2) through vigorous shaking for 5 min. After 7000 rpm centrifugation for 10 s, the filtrate was discarded. Repeat washing process several times, 10 μg of circRNA sample with 200 μL elution buffer was loaded into the column, and the mixture was shaken vigorously for 6 min. The unbound single-stranded mRNA in the effluent was collected and separated with dsRNA by 14000 × g centrifugation for 1 min. For recovery, circRNA was precipitated with LiCl.

### SEC purification of circRNA

circRNA was run through a 7.8 × 300 mm size exclusion column (Sepax Technologies; part number: 215950-7830) on AKTA FPLC (Cytiva). RNA was run in RNase-free Potassium Phosphate buffer (10 mM potassium phosphate, 1 mM EDTA, pH = 6) at a flow rate of 0.4 mL/min. For recovery, circRNA was precipitated with LiCl.

### Ultra-pure circ-eGFP production

The ultra-pure circ-eGFP was obtained according to (8). Firstly, the raw IVT-produced circ-eGFP was treated with RNase R to remove the majority of linear RNA. Then the RNase R-treated circ-eGFP was collected using LiCl precipitation and subsequently purified by SEC. The late elution peak was collected and treated with phosphatase to obtain the final product.

### Cellular fluorescence and flow cytometry

For HeLa cells, 400 ng of RNA was transfected into 4× 10^4^ cells/200 μL per well of a 48-well plate using Lipofectamine 3000 (Invitrogen) according to the manufacturer’s instructions. 24 hours after transfection, cells were imaged on a fluorescence microscope (OLYMPUS CKX53). For flow cytometry, cells washed with PBS three times and redispersed in 100 μL of PBS before being analyzed on a flow cytometer (ACEA NovoCyte, Agilent). The data were analyzed using FlowJo software V10.8. The transfection efficiency was quantified as the percentage of GFP-positive cells to the total cell populations. The expression level of GFP mRNA was represented by the mean fluorescence intensity (MFI) of fluorescein isothiocyanate (FITC) in positive cells.

### Cell transfection and RNA isolation

To obtain PMA-differentiated THP-1 cells, THP-1 cells were seeded at a density of 2.5 × 10^5^ cells/500 μL per well in a 12-well culture plate and stimulated with 100 ng/mL of PMA for 48 hours. A549 cells were seeded at a density of 2 × 10^5^ cells/500 μL per well in a 12-well culture plate 24 hours prior transfection. A total of 200 ng of RNA or 40 ng of HMW or intron fraction was transfected into A549 and THP-1 cells using Lipofectamine 3000 (Invitrogen™) according to the manufacturer’s instructions. All cells were washed with PBS once and lysed 6 hours after transfection. Total RNAs were extracted with FastPure Cell/Tissue Total RNA Isolation Kit V2 (Vazyme, China) according to the manufacturer’s protocol.

### RT-qPCR

0.1 mg of total RNAs were served as template for reverse transcription by using PrimeScript™ II 1st Strand cDNA Synthesis Kit (TAKARA) according to manufacturer’s protocols. After reverse transcription, cDNAs were transcribed with Taq Pro Universal SYBR qPCR Master Mix (Vazyme, China) according to the manufacturer’s protocol. Primer pairs were selected using the Primer Blast designing tool (https://www.ncbi.nlm.nih.gov/tools/primer-blast/). Relative expression of mRNA abundance was determined from three independent experiments and was normalized by housekeeping gene B2M.

### RNA sequencing

RNA quantity and quality assessment were performed as described previously (32). RNA-seq libraries were constructed using AHTS Universal V8 RNA-seq Library Prep Kit for Illumina (Vazyme, China) following the manufacturer’s instructions, and then sequenced on SURFSeq 5000 platform (GeneMind Biosciences LTD., China) with PE150. After filtering the adapter, ploy-N and low-quality reads, the reads were aligned to human reference genome (GRCh38) with Histat2 (33). The mapped reads were subjected to StringTie to perform expression quantification (34). Differentially expressed genes/transcripts (DEGs/DETs) were identified by DESeq2 (p.value DEGs<0.05; |log2(fold change)| ≥ 1) (35). KEGG and GO enrichment was conducted with the R package clusterProfiler (36).

## Acknowledgments

We thank Dr.Mingjie Luo and Dr. Pengwei Xu (GeneMind Biosciences Company Limited, Shenzhen, China) for their technical support with RNA sequencing.

## Author contributions

Z.Z., W.L., D.W. and Y.C. designed the experiments. Z.Z., W.L., D. L., and X.Y. J conducted the experiments and performed the data analysis. D.W. and Y.C. were responsible for research supervision, coordination, and strategy. Z.Z. and W.L. drafted the manuscript. D.W. and Y.C. reviewed and edited the manuscript. All authors reviewed and approved the final version of the manuscript.

## Data and materials availability

All data needed to evaluate the conclusions in the paper are present in the paper and/or the Supplementary Materials. Additional data related to this paper may be requested from the authors.

