## Supplemental Figures and Tables for "Mitigating Cellular Dysfunction by Addressing Contaminants in Synthetic circRNA"

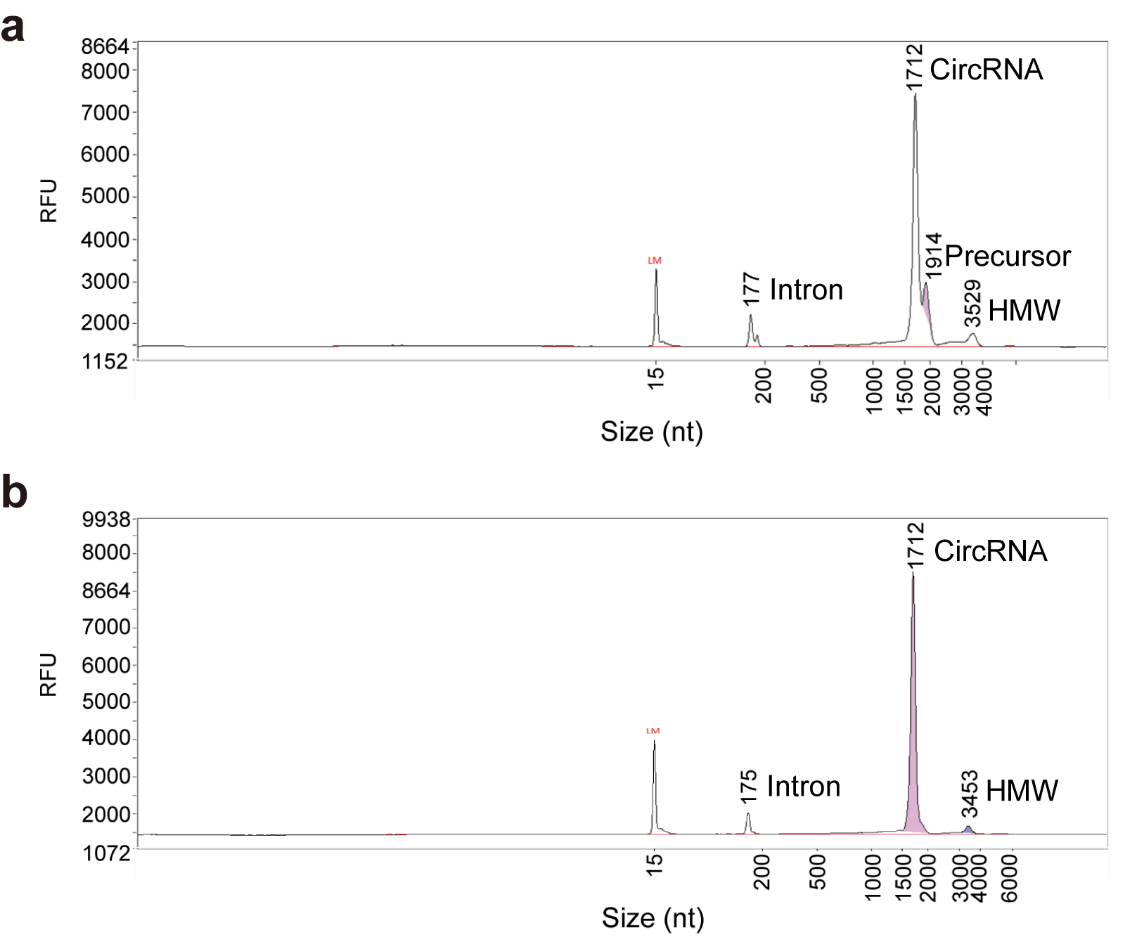

Fig. S1. Capillary electrophoresis analysis of circ-eGFP.

**a** IVT-synthesized circ-eGFP (unpurified).

**b** IVT-synthesized circ-eGFP after RNase R treatment (RR+).

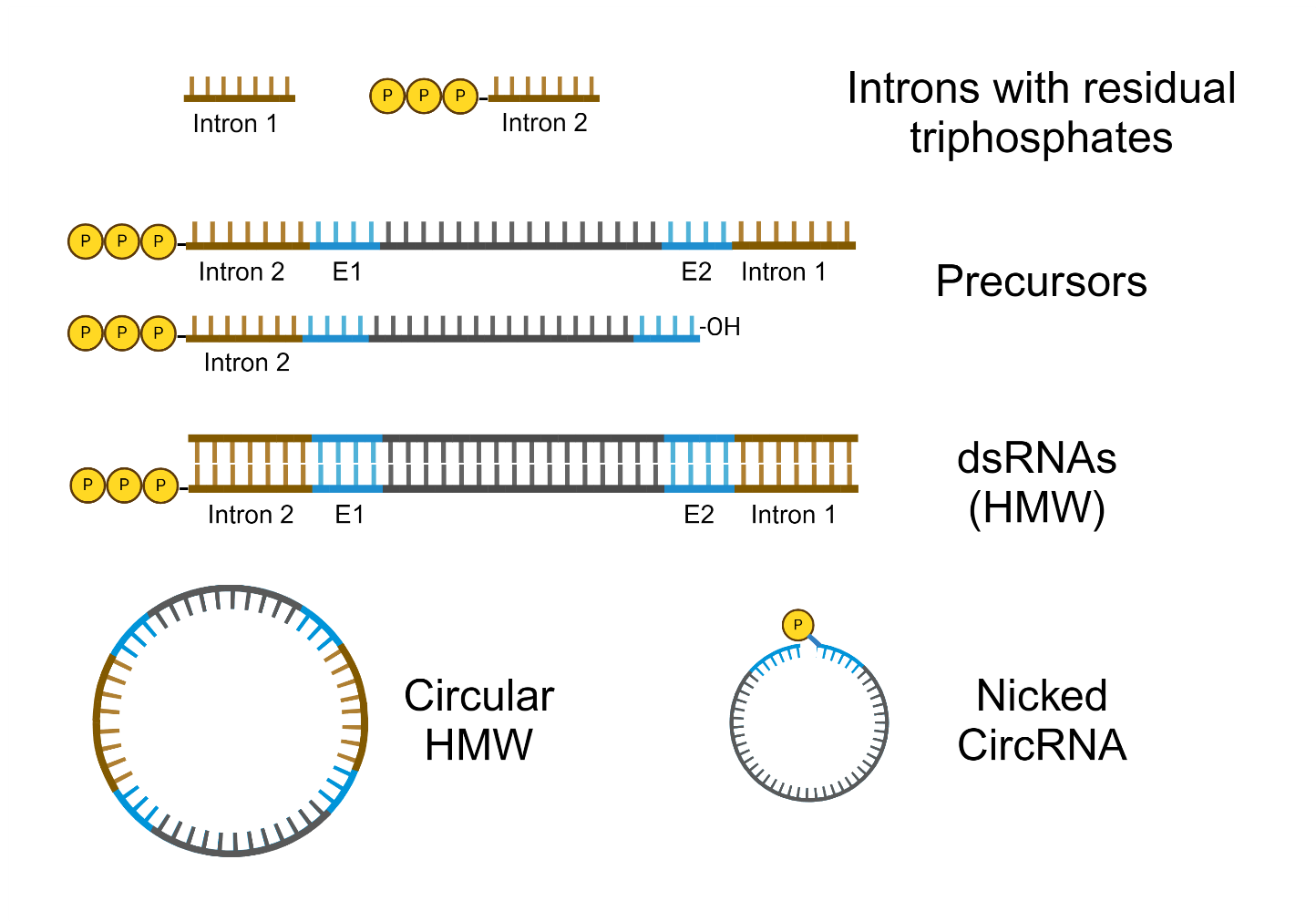

Fig. S2. The byproducts in synthetic circRNA may elicit innate immune response, including dsRNA, 5’ triphosphates introns and precursors, and hydrolyzed circRNA (nicked RNA). (Created with BioRender.com)

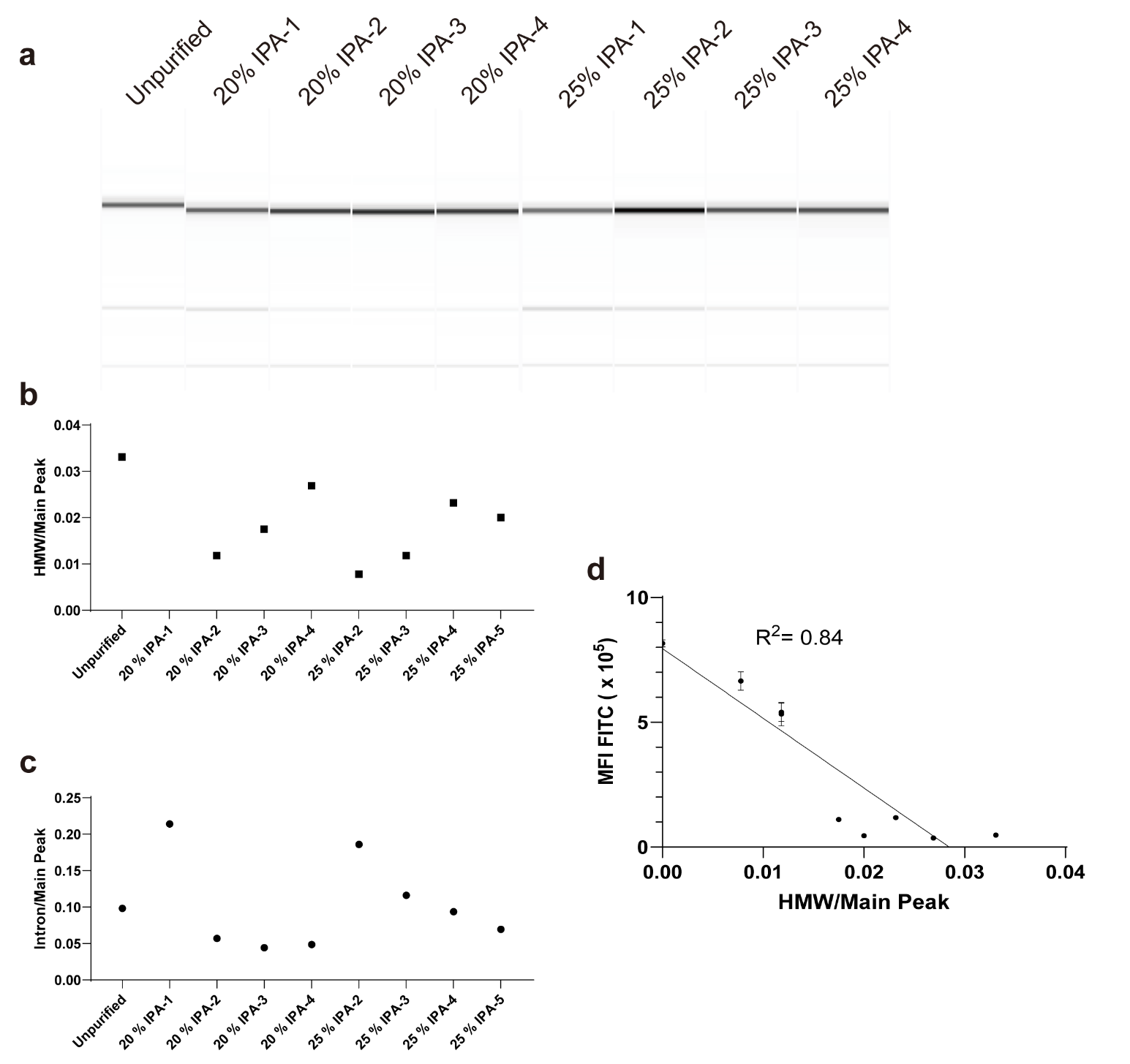

Fig. S3. Capillary electrophoresis analysis of circRNA fractions in Figure 3A.

**a** Capillary electrophoresis of circRNA fractions.

**b-c** Distribution of intron and HMW byproducts across different fractions as detailed in Figure 3A, respectively.

**d** Linear correlation between MFI and HMW ratio.

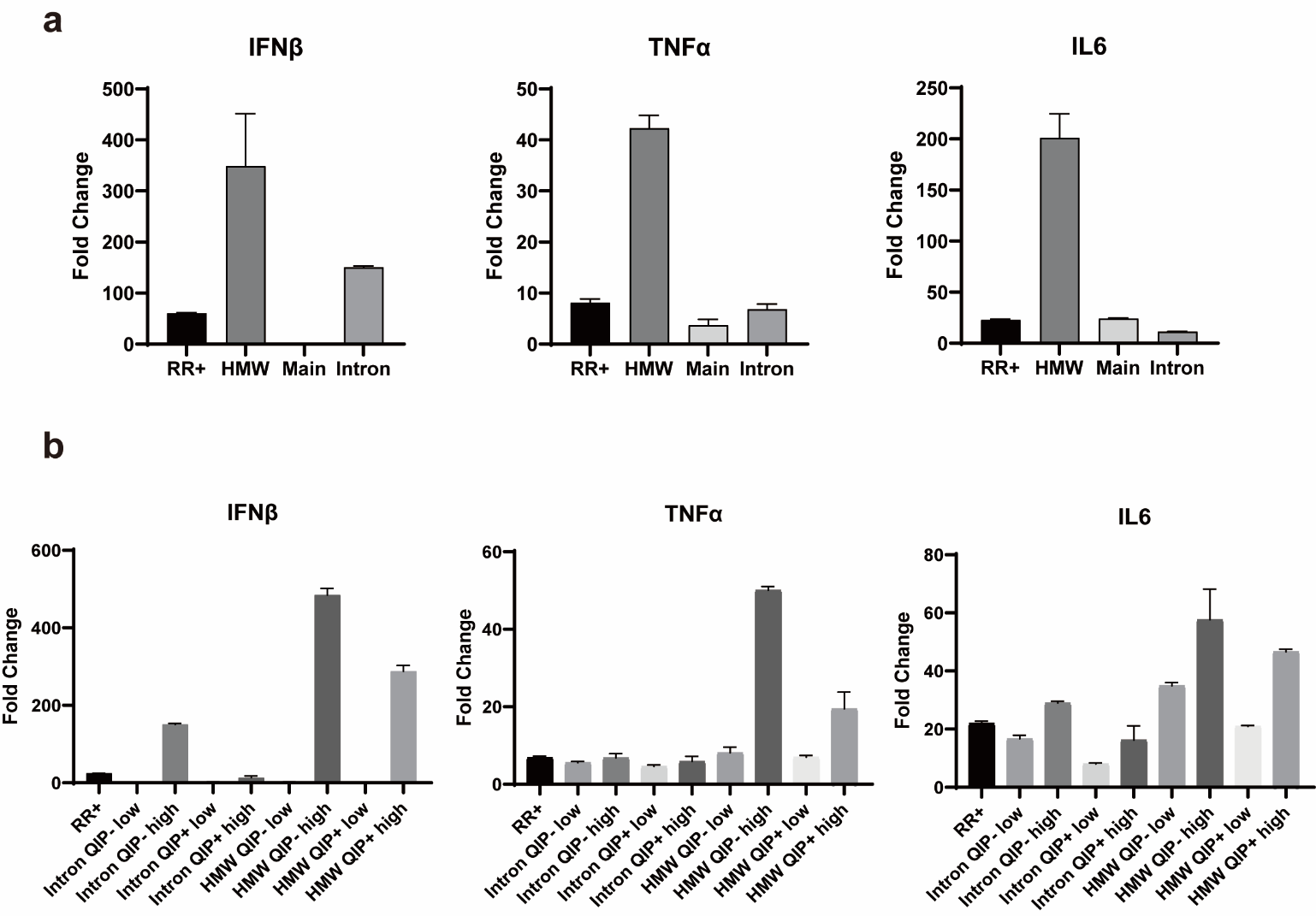

Fig. S4. Immunogenicity assessment of circ-eGFP fractions with enzymatic treatments.

**a** Induction of IL6, TNFα and IFNβ transcripts 6 hours after transfection of 40 ng of HMW or introns, 200 ng of RR+circ-eGFP, main fraction into 0.2 million A549 cells, respectively. B2M mRNA was used as housekeeping controls. Data presented as means ± standard deviations (SDs) of three biological replicates.

**b** Induction of IL6, TNFα and IFNβ transcripts 6 hours after transfection of A549 cells with the 40 ng or 200 ng of intron and HMW before and after phosphatase treatment (intron-P, HMW-P), or 200 ng of main fraction after phosphatase treatment (main-P). B2M mRNA was used as housekeeping controls. Data presented as means ± SDs of three biological replicates.

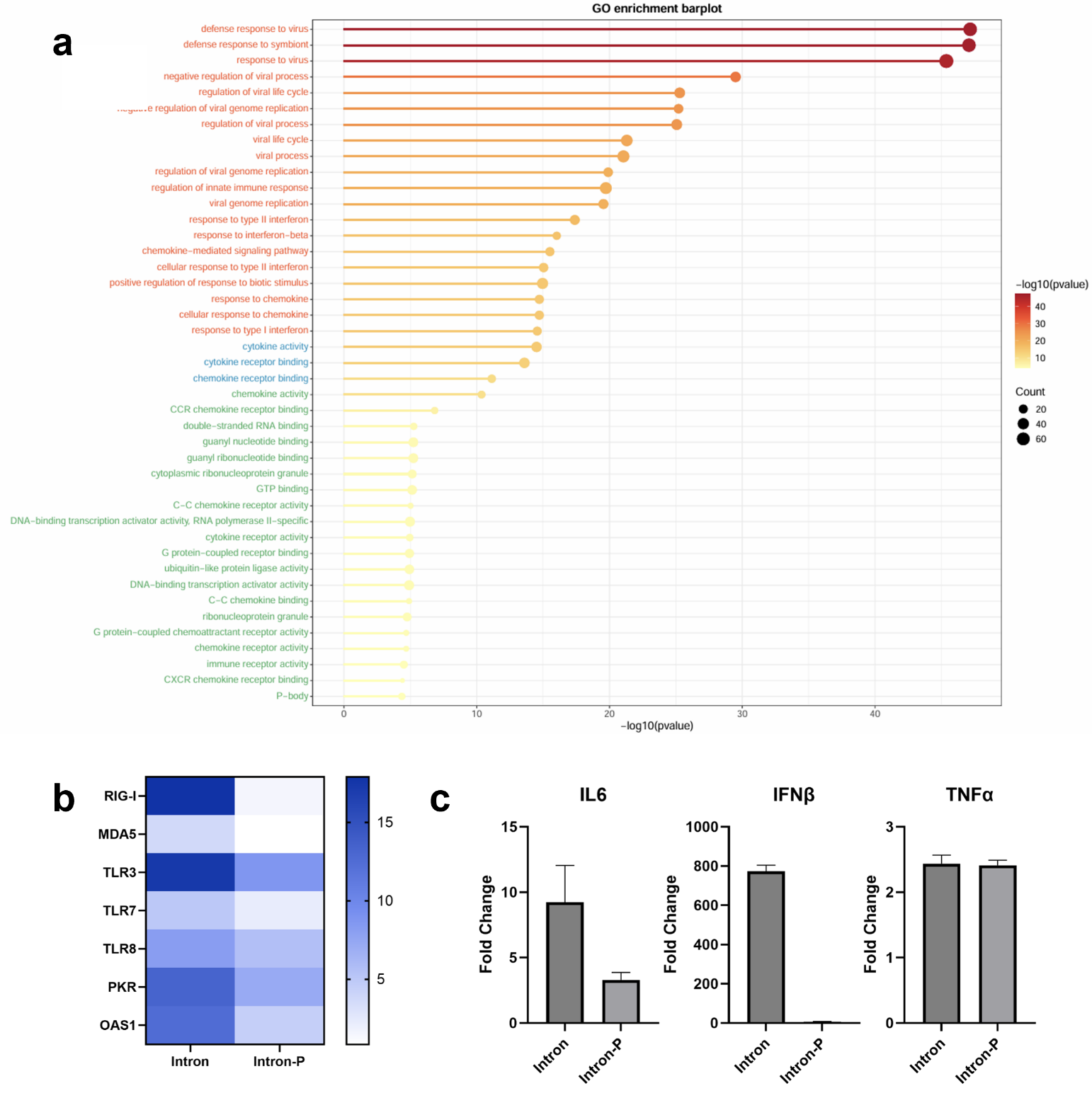

Fig. S5. Immunogenicity assessment of intron before and after phosphatase treatment.

**a** GO enrichment analysis between PMA-differentiated THP-1 cells transfected with intron after phosphatase treatment (intron-P) and intron.

**b** Induction of RIG-I, MAD5 and TLR3, TLR7 transcripts 6 hours after transfection of intron or intron-P into PMA-differentiated THP-1 cells. B2M mRNA was used as housekeeping controls. Data presented as means of three biological replicates.

**c** Induction of IL6, TNFα and IFNβ transcripts 6 hours after transfection of intron or intron-P into PMA-differentiated THP-1 cells. B2M mRNA was used as housekeeping controls. Data presented as means ± SDs of three biological replicates.

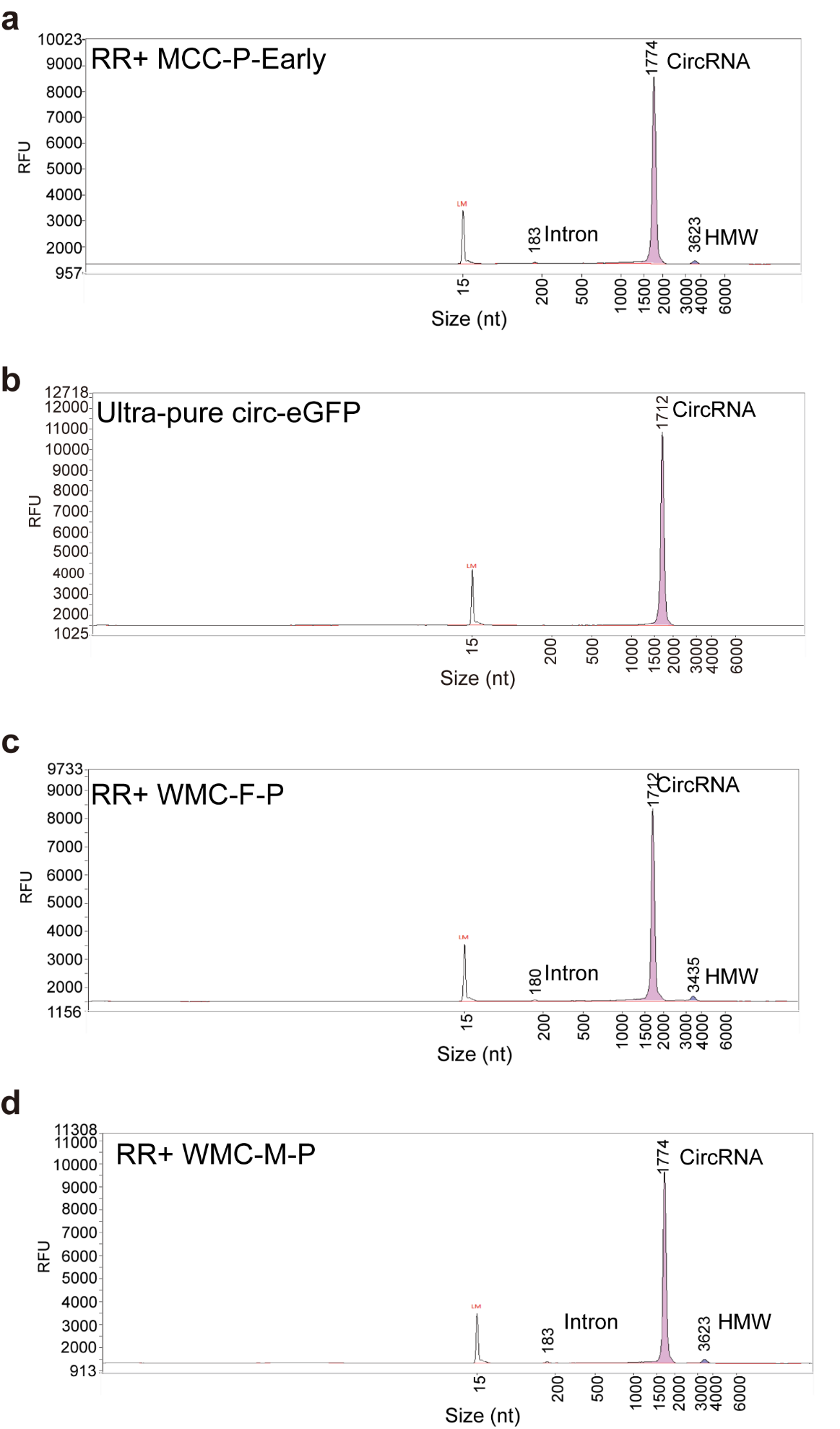

**Fig. S6. Capillary electrophoresis of RR+MCC-P, ultra-pure, WMC-F-P and RR+WMC-M-P circ-eGFP.**

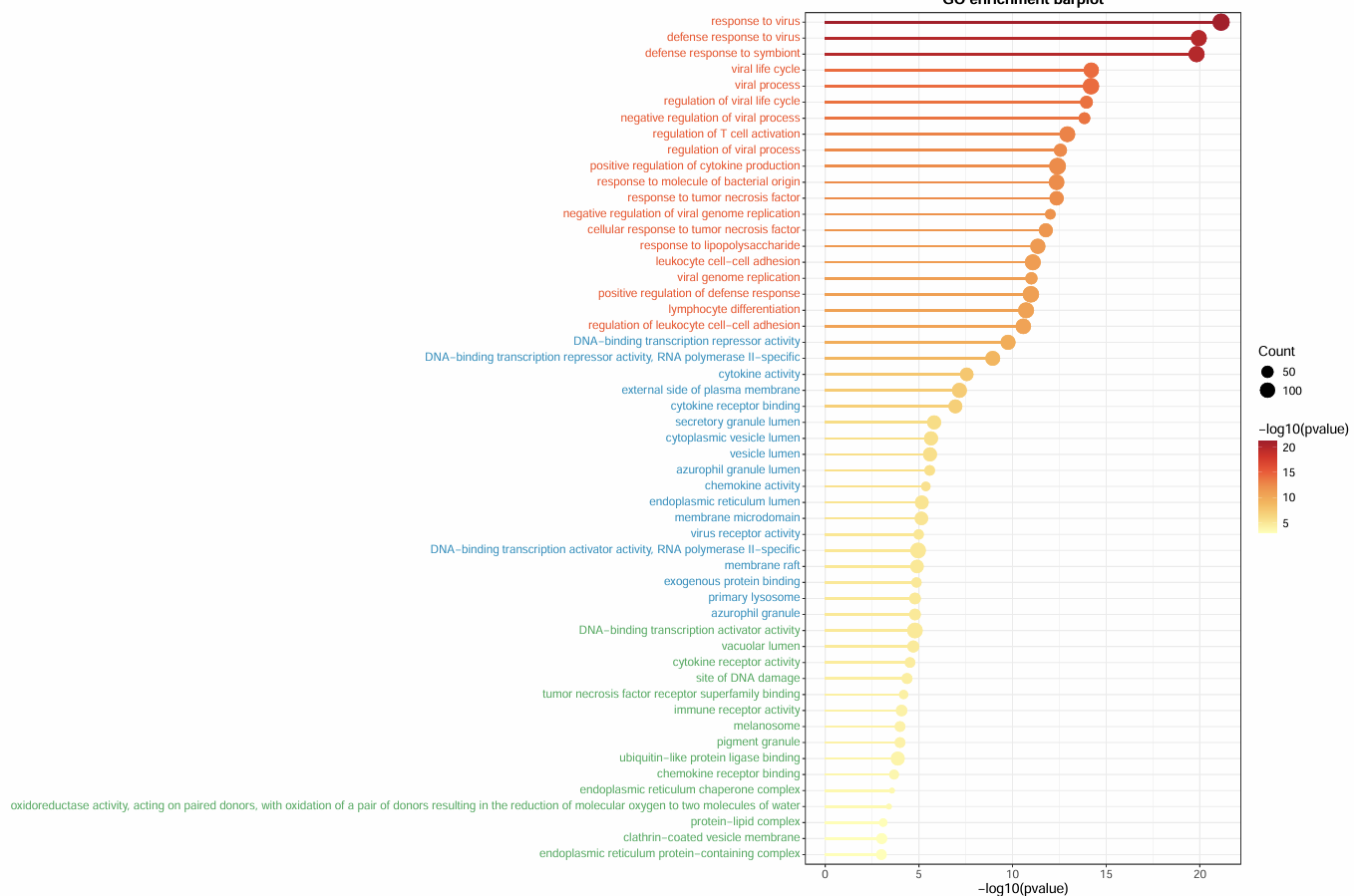

Fig. S7. GO enrichment analysis between PMA-differentiated THP-1 cells transfected HMW-MCC and HMW.

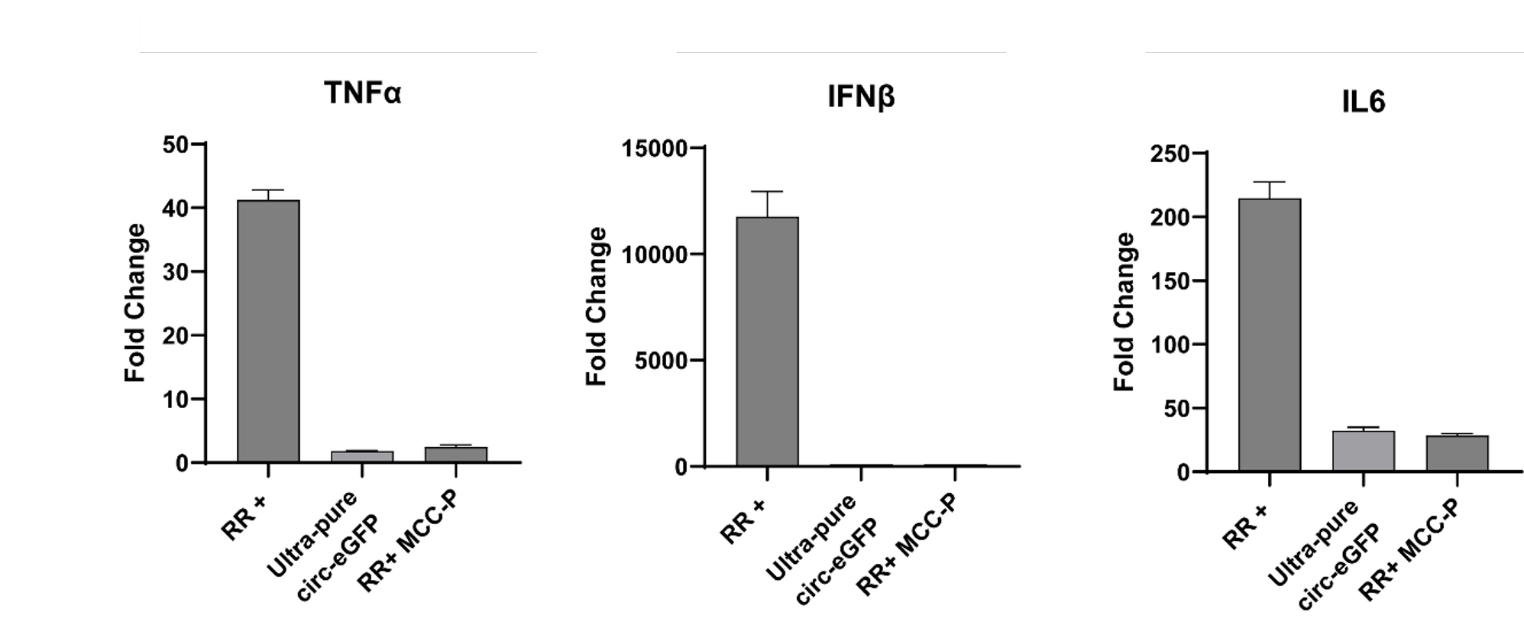

Fig. S8. Induction of IFNβ, TNFα, and IL6 transcripts 6 hours after transfection of PMA-differentiated THP-1 cells with ultra-pure, RR+MCC-P, and unpurified circ-eGFP.

B2M mRNA was used as housekeeping controls. Data presented as means ± SDs of three biological replicates.

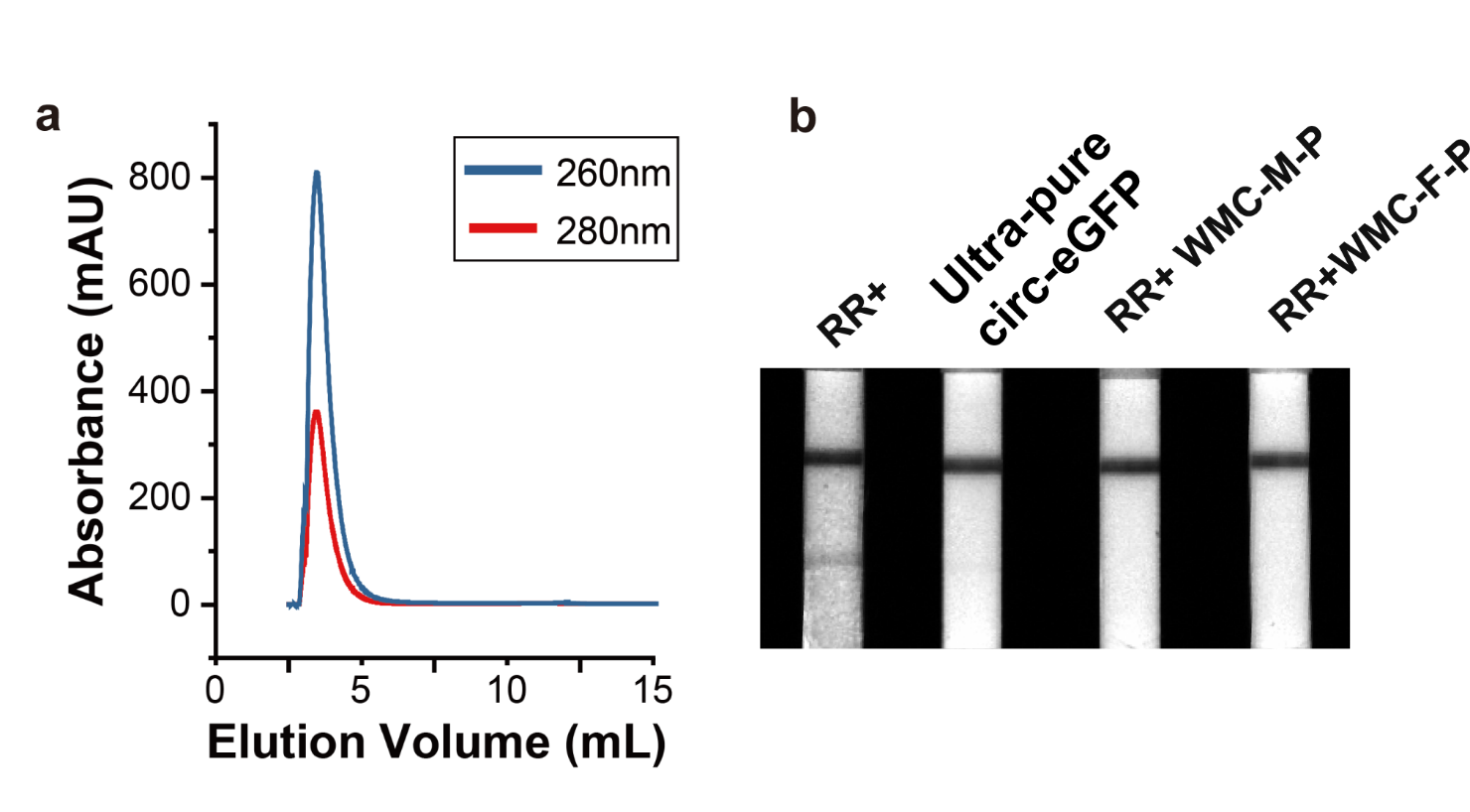

Fig. S9. WMC chromatography to purify circRNA.

a WMC separation of 200 μg of RR+circ-eGFP using FPLC.

b LFSA test of RR+, ultra-pure, RR+WMC-M-P and RR+WMC-F-P. circ-eGFP.

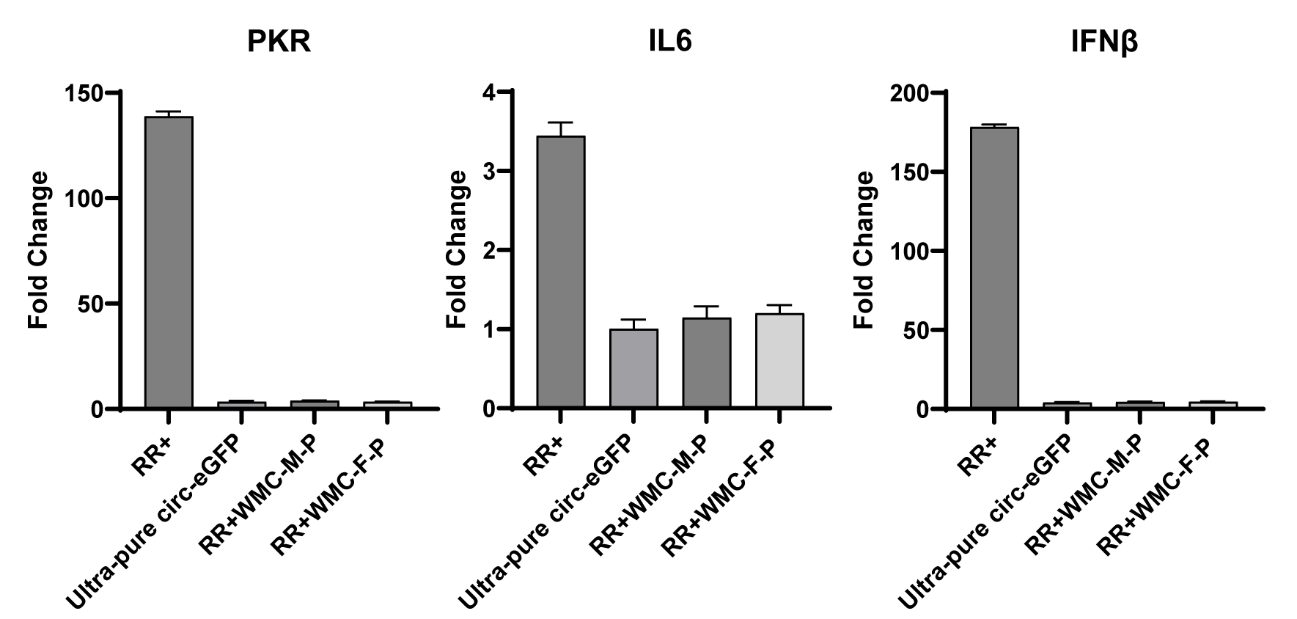

Fig. S10. Induction of PKR, IL6, and IFNβ transcripts 6 hours after transfection of PMA-differentiated THP-1 cells with RR+, ultra-pure and RR+WMC-F-P circ-eGFP.

B2M mRNA was used as housekeeping controls. Data presented as means ± SDs of three biological replicates.

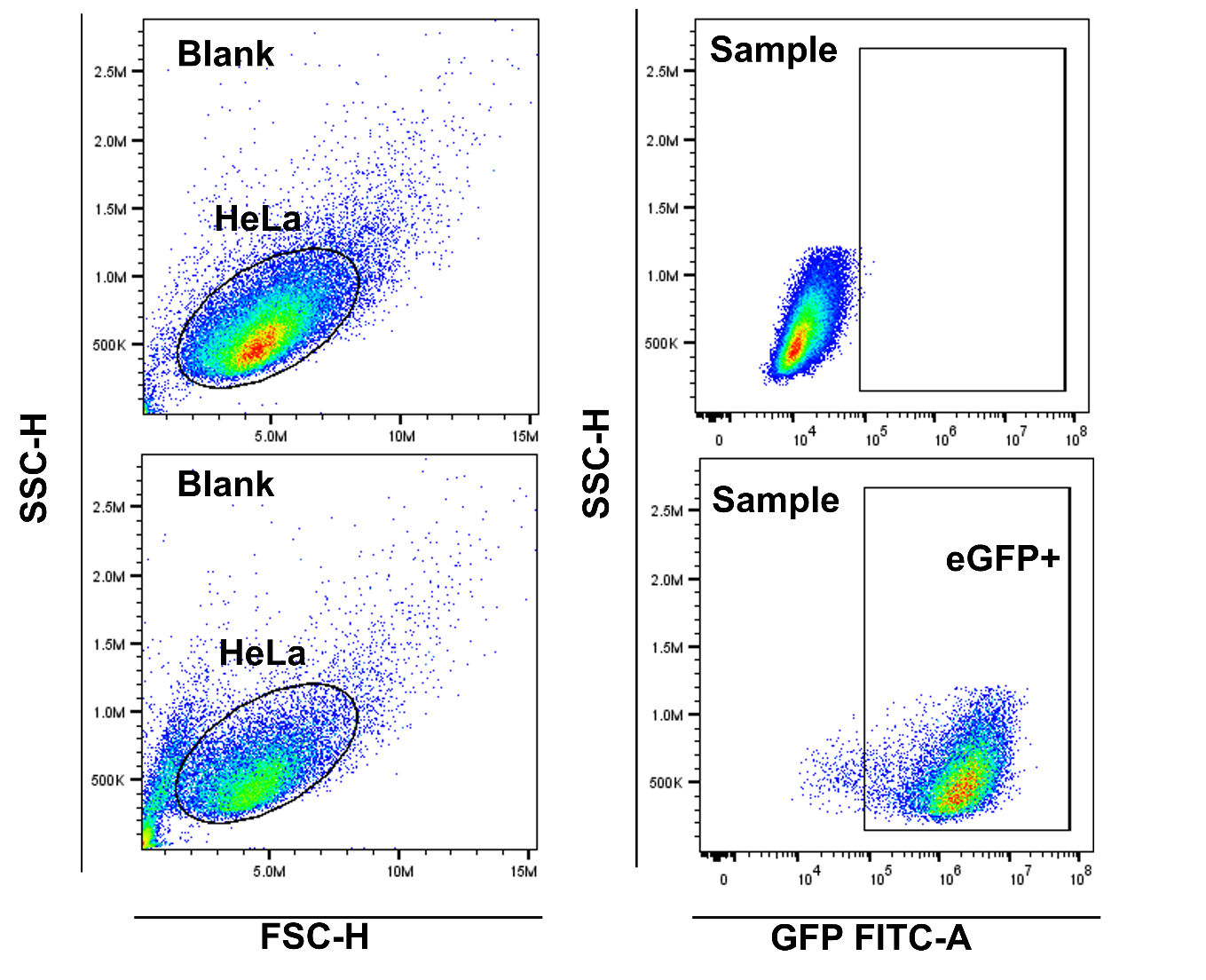

Fig. S11. Schematic and representative flow plots showing GFP positive HeLa cells .

Table S1 Intron and HMW proportions to main peak of different fractions in Figure 3A.

| **Sample** | **Intron/Main Peak** | **HMW/Main Peak** |
| --- | --- | --- |
| Unpurified | 0.098 | 0.033 |
| 20 % IPA-1 | 0.214 | Not Detected |
| 20 % IPA-2 | 0.057 | 0.012 |
| 20 % IPA-3 | 0.044 | 0.018 |
| 20 % IPA-4 | 0.049 | 0.027 |
| 25 % IPA-2 | 0.186 | 0.008 |
| 25 % IPA-3 | 0.116 | 0.012 |
| 25 % IPA-4 | 0.094 | 0.023 |
| 25 % IPA-5 | 0.070 | 0.020 |

Table S2 Reagents and resources

| **Chemicals, Peptides, and Recombinant Proteins** | **SOURCE** | **IDENTIFIER** |
| --- | --- | --- |
| PMA | Sigma | Catalog # P1585 |
| Lipo3000 | Invitrogen™ | Catalog # L3000015 |
| Human recombinant IL-2 | STEMCELL^TM^ | Catalog # 78145.1 |
| ImmunoCult™ Human CD3/CD28 T Cell Activator | STEMCELL^TM^ | Catalog # 17951 |
| ImmunoCult™-XF T Cell Expansion Medium | STEMCELL^TM^ | Catalog # 10981 |
| RNase R | Epicentre | Catalog # RNR07250 |
| DNase I | Vazyme | Catalog # EN402-01 |
| Quick CIP | New England Biolabs | Catalog # M0525V |
| HEPES | Solarbio | Catalog # 7365-45-9 |
| NaCl | Solarbio | Catalog # 7647-14-5 |
| EDTA | Thermo Fisher | Catalog # J15694 |
| IPA | Innochem | Catalog # I1700 |
| Potassium Phosphate Buffer | Bioleaper | Catalog # BR4000306 |
| **Critical Commercial Assays** |  |  |
| Neon™ Transfection System 10 μL Kit | Invitrogen™ | Catalog # MPK1096 |
| T7 High Yield RNA Transcription Kit | Vazyme | Catalog # TR101-01 |
| FastPure Cell/Tissue Total RNA Isolation Kit V2 | Vazyme | Catalog # RC112-01 |
| PrimeScript™ II 1st Strand cDNA Synthesis Kit | TAKARA | Catalog # 6210A |
| Taq Pro Universal SYBR qPCR Master Mix | Vazyme | Catalog # Q712-02 |

**Table S3 Primers**

| **Primers** |  |
| --- | --- |
| B2M-F | AGGACTGGTCTTTCTATCTC |
| B2M-R | TTCATCCAATCCAAATGCGG |
| IL6-F | GACTGCAGGAACTCCTTAAAGC |
| IL6-R | GAGTAGTGAGGAACAAGCCAGAG |
| IFNβ-F | TTGAATGGGAGGCTTGAATACT |
| IFNβ-R | TAGCCAGGAGGTTCTCAACAAT |
| TNFα-F | AAGACCACCACTTCGAAACCT |
| TNFα-R | AGGCCTAAGGTCCACTTGTGT |
| TLR7-F | TGCTGTGTGGTTTGTCTGGTG |
| TLR7-R | CTCCTGGCCCCACACAAGT |
| TLR8-F | TCCGCACTTGAAACTAAGACC |
| TLR8-R | CTGGCACAAATGACATCTAC |
| OAS1-F | TGGATTCTGCTGGTGAGACC |
| OAS1-R | ATGGCCTTTGGCAAGAGGTAAG |
| RIG-I-F | AAATCAGAACACAGGCAGAGGAA |
| RIG-I-R | GTCCCATGTCTGAAGGCGTAA |
| MDA5-F | GCATATGCGCTTTCCCAGTG |
| MDA5-R | CTCTCATCAGCTCTGGCTCG |
| PKR-F | TCCATGGGGAATTACATAGGC |
| PKR-R | AGCGGCCAATTGTTTTGCTT |
| TLR3-F | CCTTTTGCCCTTTGGGATGC |
| LR3-R | TGAAGTTGGCGGCTGGTAAT |
